# Conserved inhibitory mechanisms of root-associated bacteria towards *Arabidopsis thaliana* and *Chlamydomonas reinhardtii*

**DOI:** 10.64898/2026.09.14.751450

**Authors:** Tzila Vikinyanski, Elad Meilin, Andrei Makhon, Tingting Song, Moshe Alon, Noémie Alon Cudkowicz, Samiksha Saxena, Avital Akerman-Arad, Ola Karmi, Shaul Pollak, Asaf Levy, Manuel Kleiner, Michal Breker, Omri M. Finkel

## Abstract

Soil bacteria colonize plants without causing visible disease, yet many suppress host growth. The prevalence and evolutionary reach of this cryptic inhibition are unknown, because phenotyping hundreds of isolates in plants is prohibitively slow. Here we use the unicellular algae *Chlamydomonas reinhardtii* as a scalable proxy for the green lineage. We screened 148 plant-associated bacteria on Chlamydomonas lawns. Seven of eight Chlamydomonas inhibitors also inhibited the vascular model plant *Arabidopsis thaliana*. In one of these dual inhibitors, *Burkholderia sola* MF6, transposon screening and proteomics implicated type VI secretion, tight-adherence pili and secreted metallopeptidases. Over half the non-inhibitory mutants also failed against Arabidopsis, indicating shared mechanisms towards plants and algae. In the algae, infection triggers rapid deflagellation, then non-lytic regulated cell death. The algal zinc/iron transporter ZIP3 promotes inhibition, whereas the cathepsin X protease CEP12 restrains it, and ZIP3 is epistatic to CEP12. Cryptic inhibition is thus ancient and widespread, and a unicellular algae uncovers it at scale.

---

Plants interact with diverse microorganisms, collectively termed the plant microbiota^1^. Members of the microbiota evolved to manipulate the plant to become more habitable to them^2–5^. Many of these microbes reciprocate mutualistically, providing nutritional or protective benefits^6,7^. But often, plant-microbe interactions are costly for the plant and compromise its growth^4^. This is apparent in acute disease, but plants also endure “chronic infections” carrying subtle, difficult to diagnose costs^8^. Unlike acute pathogens that cause symptoms like visible necrosis^9^, these cryptic antagonists are found in seemingly asymptomatic plants, and only appear to be plant growth inhibitors when inoculated into gnotobiotic systems^4,8,10^, an observation reinforcing the microbiota’s role in protecting from such antagonists. Despite this protection, the cumulative cost of multiple chronic interactions with antagonistic bacteria may be substantial, explaining why plants do not always reach their growth potential, even when no disease is apparent^11^. Similarly to opportunistic pathogens^12,13^, potentially harmful microbes found within seemingly asymptomatic contexts need to be monitored and understood.

To optimize agricultural yields, we must understand both the magnitude and mechanisms of cryptic antagonism. But screening for microbially-induced phenotypes in multicellular plants is restricted by long life cycles, physical space, and biological variability. This phenotyping bottleneck prevents the high-throughput analysis necessary to screen thousands of microbial taxa in soil, and identify virulence factors and protective agents. Bridging this gap requires a model system that combines scalability and relevance.

The unicellular green algae *Chlamydomonas reinhardtii* is a canonical model for plant cell biology^14^. It shares a deep evolutionary history with land plants, retaining extensive homology in gene families governing photosynthesis, metabolism, and stress responses^15^. Chlamydomonas was recently proposed as a model for plant microbiome research, given that the algal phycosphere recruits similar taxa to the Arabidopsis rhizosphere^16^. However, it remains to be determined whether overlap in community structure translates to conservation of function and whether bacteria utilize analogous mechanisms to inhibit both algae and plants. If this is the case, Chlamydomonas could serve as a proxy for rapid, high-throughput discovery of plant-antagonistic mechanisms, and provide the opportunity to study host genetics systematically.

Here, we investigated evolutionary conservation of bacterial inhibition toward photosynthetic eukaryotes. By screening a collection of bacteria isolated from Arabidopsis roots, we observed that inhibition towards Chlamydomonas and Arabidopsis overlaps. Shared inhibitory strains spanned three bacterial phyla. Focusing on the dual inhibitor *Burkholderia sola* MF6 (*Bso* MF6). Using genetic screens, we identified multiple bacterial genes required for inhibiting both plants and algae, indicating a shared virulence strategy. Furthermore, we discovered that the Chlamydomonas zinc/iron transporter *ZIP3* confers susceptibility to *Bso* MF6 and that the cysteine endopeptidase *CEP12* confers resistance. These results reveal conserved physiological vulnerabilities, such as metal homeostasis, that soil bacteria exploit.

Unicellular algae therefore offer a rapid, reliable, and evolutionarily relevant model system to decode mechanisms of plant growth inhibition at scale.

## Results

### Overlapping inhibition among diverse bacteria towards Chlamydomonas and Arabidopsis

To assess the overlap in bacterial inhibition towards plants and algae, we performed an algal inhibition (halo) assay using 148 bacterial strains isolated from Arabidopsis roots, spanning four phyla (Fig. 1a, b, Extended Data Figs. 1a-c Supplementary Table S1)^6^. Eight strains inhibited Chlamydomonas growth (Fig. 1a, b and Extended Data Fig.1c, Supplementary Tables S1 and S2): two *Burkholderia*, four *Paenibacillus*, and two *Pseudomonas* (an additional *Pseudomonas* strain, MF395, also inhibited Chlamydomonas, but since no data is available on its effect on Arabidopsis, it is not counted). Seven of these eight are also plant root growth inhibitors (Fig.1b)^4^.

**Figure 1.**
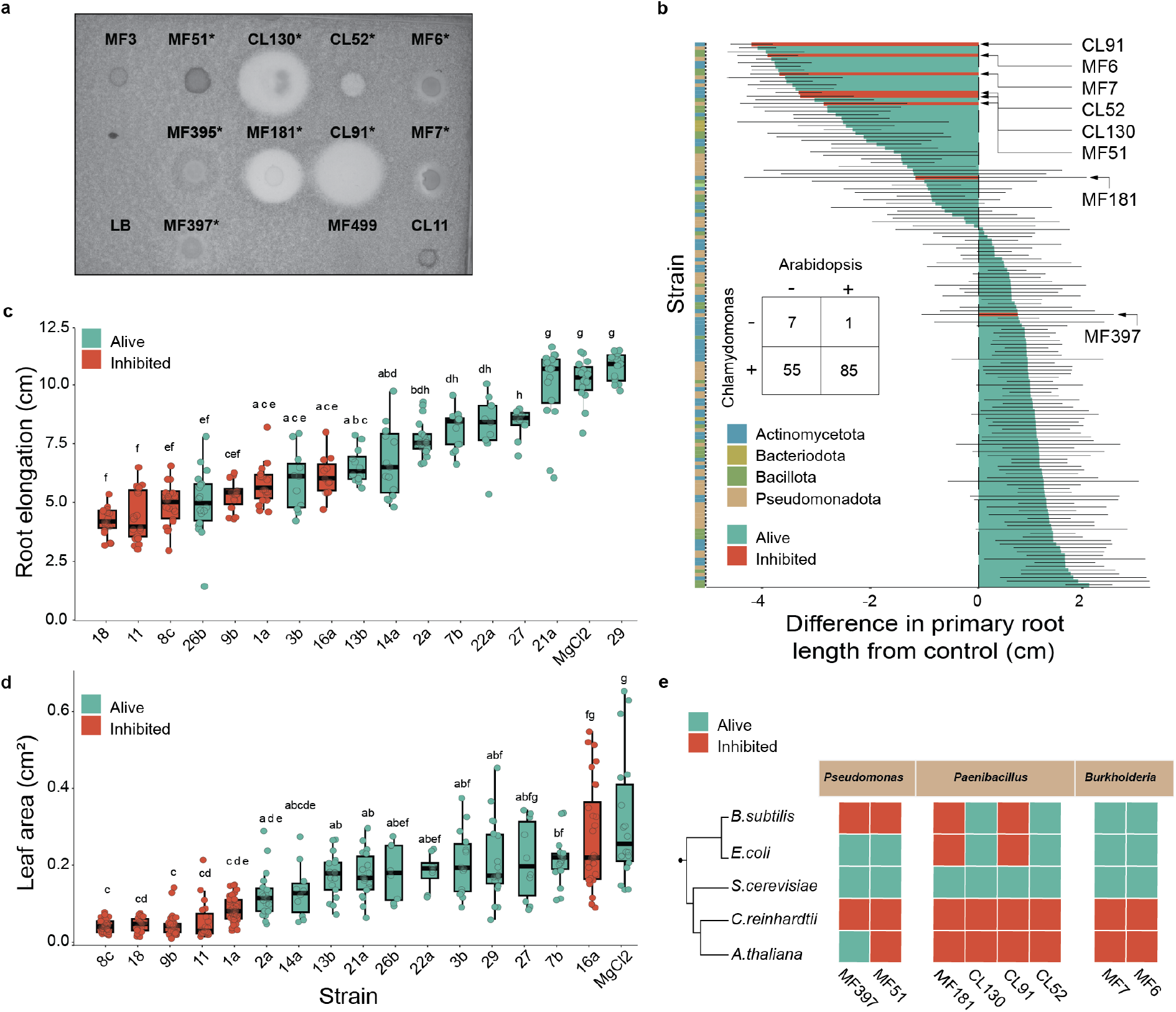
Bacterial inhibition toward Chlamydomonas predicts inhibition toward Arabidopsis. (a) Representative halo assay. Overnight cultures of Arabidopsis root-derived bacterial strains were spotted (4 µl, OD = 1) onto a Chlamydomonas lawn embedded in TAP soft agar and imaged after 3 days of continuous light at 21 °C. Colorless zones surrounding a colony indicate inhibition of algal growth; LB alone served as the negative control. Marked with an asterisk are considered inhibitors. (b) Effect of 148 Arabidopsis root-derived bacterial strains on Arabidopsis primary root elongation, expressed as the difference from uninoculated control (cm) and ranked from most to least inhibitory (data from Finkel et al)^4^. Bars are colored by each strain’s effect on Chlamydomonas in the halo assay in (a): red, halo-forming (inhibited); green, no halo (not-inhibited). The color strip on the left denotes bacterial phylum. Arrows indicate the strains that inhibited Chlamydomonas. Statistical significance (Welch’s *t*-test) is reported in Table S2. 22 of the 148 strains score 0 for growth on TAP alone. Inset: overlap between inhibition of Chlamydomonas and of Arabidopsis among 148 root isolates with both measurements. Chlamydomonas inhibition was scored by halo assay; Arabidopsis root growth inhibition is calculated from Finkel et al.^4^ Bacteria that inhibited Chlamydomonas were more likely to inhibit Arabidopsis than the collection as a whole (7/8 versus 62/148; odds ratio 10.8, *p* = 0.0097, Fisher’s exact test). (c and d) Primary root elongation (c) and leaf area (d) of Arabidopsis seedlings 12 days after transfer onto plates inoculated with bacteria isolated from pine-forest soil using Chlamydomonas halo formation as the selection criterion. Strains are ordered by increasing value; MgCl₂ denotes the uninoculated control. Colors indicate Chlamydomonas inhibition status as in (b). Boxes show the median and interquartile range, whiskers extend to 1.5× IQR, and points represent individual seedlings. Groups not sharing a letter differ significantly (*p* < 0.05, Tukey’s HSD following one-way ANOVA (e) Target range of the Chlamydomonas-inhibitory strains across the tree of life. Each strain (columns, grouped by genus) was spotted onto lawns of *Bacillus subtilis*, *Escherichia coli*, *Saccharomyces cerevisiae*, and Chlamydomonas, and inoculated onto Arabidopsis seedlings. Red, growth inhibited; green, not inhibited. The cladogram on the left indicates the phylogenetic relationships among target organisms. MF3, MF395, MF499, CL11 were excluded. See also Extended Data Fig. 1 and Supplementary Tables S1-S5.

To test if Chlamydomonas inhibition reliably indicates plant inhibition beyond this isolate collection, we enriched plant-antagonistic bacteria from soil using Chlamydomonas inhibition as a readout. We isolated bacterial colonies from halos formed on Chlamydomonas lawns exposed to forest soil slurries (Extended Data Fig. 1d, e). Six of the 16 bacteria isolated this way formed halos on Chlamydomonas lawns (Extended Data Fig. 1e). All six halo-forming bacteria, two *Pseudomonas*, one *Paenibacillus*, and one *Streptomyces,* also inhibited Arabidopsis root growth relative to an uninoculated control (Fig. 1c, Supplementary Table S1), and five also reduced shoots (Fig. 1d, Supplementary Tables S3, S4 and S5). Thus, a bacterium that inhibits Chlamydomonas is likely to inhibit Arabidopsis, independently of isolation source.

Finally, we asked whether dual inhibitors were specific to photosynthetic organisms or instead acted broadly. We tested eight dual inhibitors against Gram^-^ (*Escherichia coli*) and gram^+^ (*Bacillus subtilis*) bacteria and baker’s yeast (*Saccharomyces cerevisiae*). The eight strains showed variable ranges, from broad-spectrum inhibition (*Paenibacillus* MF181) to apparent specificity towards photosynthetic organisms (two *Burkholderia* strains and two *Pseudomonas*), suggesting that algae and land plants share susceptibility features (Fig. 1e and Extended Data Fig. 1f). Taken together, we revealed common inhibitory activity among diverse soil-dwelling bacteria towards both plants and algae, indicating that Chlamydomonas can be a useful model for studying bacterial plant inhibition.

### *Bso* MF6 inhibits Chlamydomonas partly by a contact-dependent mechanism

Bacterial inhibition of eukaryotes is often mediated by secretion systems^17^. We therefore searched the genomes of dual inhibitors for secretion systems (Fig. 2a, Supplementary Table S6). The only system common to all strains were type IVc (Tad) pili that allow tight adherence^18^. *Paenibacillus* harbored type VII secretion systems (T7SS), which mediate protein export in Gram-positive bacteria and in some species secrete effectors^19^. By contrast, some *Pseudomonas* and all *Burkholderia* encoded the contact-dependent type III (T3SS) and type VI (T6SS) secretion systems, alongside type II secretion systems (T2SS), which release proteins into the extracellular environment^20^ (Fig. 2a).

**Figure 2.**
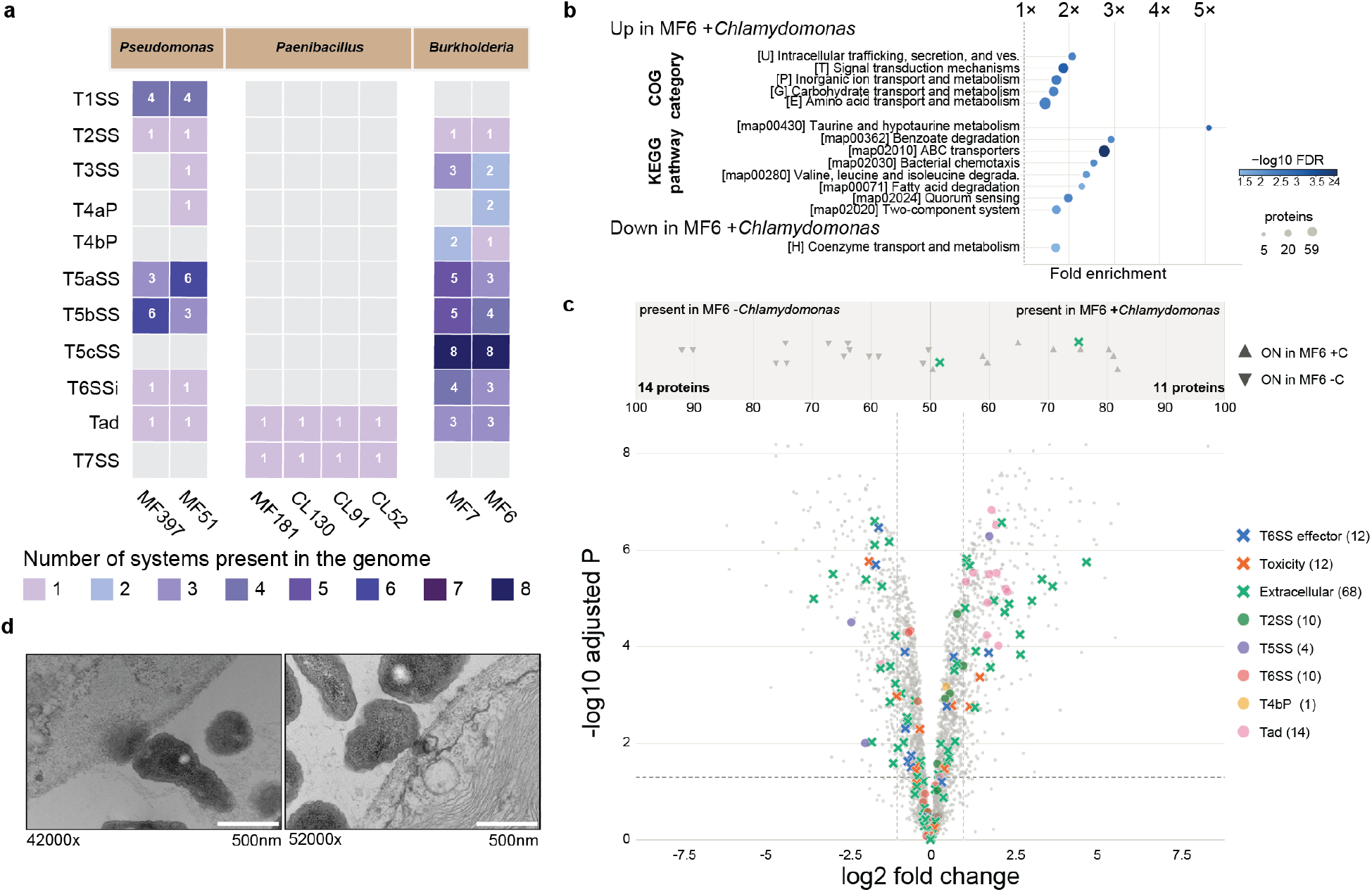
*Bso* MF6 encodes an expanded secretion-system repertoire and induces secretion-associated proteins upon contact with Chlamydomonas. (a) Secretion systems and pili identified in the genomes of the Chlamydomonas-inhibiting strains. Genomes were searched for secretion-system-associated loci; color indicates the number of complete systems of each class per genome (see Online Methods for cluster-definition criteria). Strains are grouped by genus. (b) COG categories and KEGG pathways over-represented among proteins significantly more abundant in *Bso* MF6 during co-culture with Chlamydomonas. Fold enrichment is shown on the x-axis; point size indicates the number of proteins per term and color the −log₁₀ FDR (Fisher’s exact test with Benjamini–Hochberg correction, tested against the detected bacterial proteome). (c) Differential protein abundance in *Bso* MF6 after 2 days of co-culture with Chlamydomonas versus monoculture. The volcano plot shows log₂ fold change against −log₁₀ adjusted *p*; dashed lines mark |log₂FC| = 1 and the significance threshold. Colored symbols denote proteins assigned to secretion and toxicity classes (number of proteins per class in parentheses); grey points are all other detected proteins. The upper strip shows presence/absence (“on/off”) calls, proteins detected in ≥3/4 replicates of one condition and 0-1 replicates of the other, positioned by their abundance percentile within the group in which they were detected. Downward triangles, present only in monoculture (14 proteins); upward triangles, present only in co-culture (11 proteins). (d) Transmission electron micrographs of *Bso* MF6 in contact with Chlamydomonas, 8 h after inoculation of liquid co-cultures. Bacterial cells are apposed to the algal surface, with local distortion of the algal cell wall and membrane at contact sites. Left, 42,000×; right, 52,000×; accelerating voltage 120 kV. Scale bars, 500 nm. See also Extended Data Figs. 2 and 3 and Supplementary Tables S6-S11.

To test whether algal inhibition requires contact, we separated the bacterial culture from the Chlamydomonas lawn with a micropore filter (Extended Data Fig. 2a). *Paenibacillus* inhibited through the filter, indicating a diffusible agent. *Pseudomonas* inhibition was abolished by the filter. *Burkholderia* showed an intermediate phenotype, producing smaller halos through the filter than in direct contact, indicating that inhibition is partly contact-dependent.

The apparent green-lineage specificity of *Burkholderia*, together with the large number of secretion systems they harbor (Fig. 1e), suggested a targeted mechanism rather than a generic effect such as medium acidification^21^. We therefore focused on *Burkholderia sola* MF6 (*Bso* MF6). *B. sola* is a recently described species^22^ within the *Burkholderia cepacia* complex, a clade distinguished from other *Burkholderia* by a third chromosome of approximately 1 Mbp (Extended Data Fig. 2b, Supplementary Table S7). Members of the complex are common in the rhizosphere and are also opportunistic human pathogens^23^. Unlike their close relatives, most of which are clinical isolates, all *B. sola* isolates, including MF6 and MF7, are plant-associated, and they share high sequence similarity despite originating from different sites (Extended Data Fig. 2b).

Proteomic analysis of liquid algal-bacterial co-cultures showed that *Bso* MF6 accounted for 88.8%±2.1 of unique spectra within two days. We identified 434 bacterial proteins that were enriched in the presence of Chlamydomonas (log2FC > 1, *q* < 0.05; Supplementary Tables S8, S9 and S10). The enriched KEGG and COG categories (Fig. 2b) reflect a sensing, transport and catabolism consistent with foraging for compounds released by dying algae. Among the enriched proteins were 11 of 14 detected Tad components (Fig. 2c, Supplementary Table S11), 20 proteins predicted as extracellular and one predicted T6SS effector homologous to catechol 1,2-dioxygenase (machine learning prediction; see Methods). T6SS components were not enriched, but one of the genome’s three T6SS clusters was constitutively abundant (Supplementary Tables S8 and S9). The *Bso* MF6 genome also encodes several biosynthetic gene clusters (BGCs), some potentially algicidal^24^ (Supplementary Tables S8 and S9); of these, only one, annotated as a terpene BGC, was induced.

We used transmission electron microscopy (TEM) to visualize the interaction. Within 8 h of inoculation, *Bso* MF6 cells were in direct contact with Chlamydomonas. At several contact sites, the algal cell wall and membrane appeared locally distorted, with electron-dense material continuous between the two cells (Fig. 2d). The enrichment of Tad components and the predicted T6SS effector in the presence of Chlamydomonas, together with the direct contact seen by TEM, and the reduced inhibition across a physical barrier, indicate that inhibition by *Bso* MF6 is largely contact dependent, and implicate Tad-mediated attachment in the mechanism.

### A multi-pronged attack of *Bso* MF6 towards Chlamydomonas

To identify bacterial genes required for *Bso* MF6 inhibition of Chlamydomonas, we screened a *Bso* MF6 transposon mutant library for loss of inhibition in both solid and liquid formats (Extended Data Fig. 3a, b). Of the 5,023 mutants, 187 exhibited reduced inhibition compared to wild-type *Bso* MF6, while retaining growth in TAP medium alone (Extended Data Fig. 3c, d). Secondary validation prioritized 14 non-inhibitory insertion mutants (Fig. 3a and Extended Data Fig. 3e). Ten of these had lost chromosome 3 entirely, a phenomenon previously associated with virulence loss in related *Burkholderia*^25^, suggesting that chromosome 3 contributes to *Bso* MF6 inhibition towards Chlamydomonas. Interestingly, all isolates that lost chromosome 3 had insertions within or adjacent to T6SS-associated loci (Extended Data Fig. 3f, Supplementary Tables S12 and S13). Four of these, 27D6, 51B12, 54B7 and 57A4, carried integrations within or immediately adjacent to an ATP-dependent OLD-family endonuclease gene, a family recurrently deployed as a polymorphic toxin payload in Rhs and T6SS^26^, located 2 kb upstream of a *vgrG* gene and 15 kb from a T6SS structural operon (Fig. 3a, Supplementary Table S12). The convergence of four independent insertions strongly implicates this gene. The four additional T6SS-related insertions include 42E1 that maps 83 bp upstream of the amidase immunity gene *tai4*; 34F7 that falls between a hypothetical ORF and a divergently transcribed *vgr* gene; 24G11 that lies upstream of a LysR-family regulator divergently oriented to a T6SS effector cassette; and 50A5 that lies 49 bp downstream of a Tae4-family amidase effector gene. Every one of these mutants had also lost chromosome 3, suggesting a possible causal link between loss of T6SS activity and loss of the chromosome. Our data cannot establish whether either is sufficient for loss of inhibition alone.

**Figure 3.**
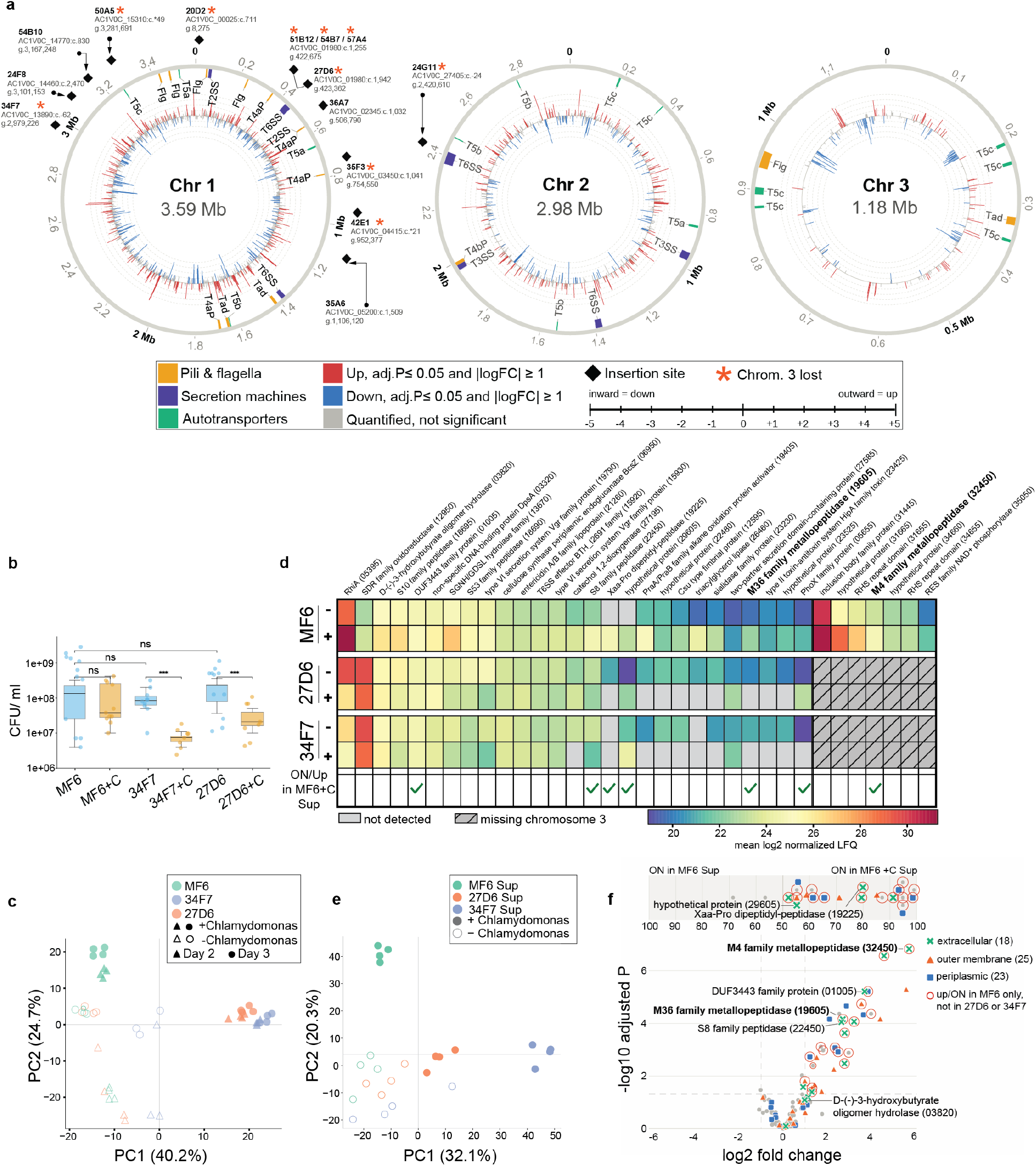
Genetic and proteomic analysis of non-inhibitory *Bso* MF6 mutants identifies candidate proteins induced by Chlamydomonas. (a) Circular genome maps of the three *Bso* MF6 chromosomes showing transposon insertion sites and the proteomic response to Chlamydomonas. Outer ring, secretion systems (purple), pili and flagella (orange), and autotransporters (green). Inner rings, differential protein abundance in co-culture with Chlamydomonas versus monoculture: bars extend outward for increased and inward for decreased abundance (log₂ fold change, scale −5 to +5); red, significantly increased; blue, significantly decreased (*q* ≤ 0.05, |log₂FC| ≥ 1); grey, quantified but not significant. Diamonds mark the insertion sites of the non-inhibitory transposon mutants, labeled with mutant name, disrupted locus and genomic coordinate. In cases of intergenic insertion, the nearest ORF is reported. A dash in the insertion coordinate denotes 3’ position and an asterisk denotes 5’ position. Orange asterisks denote mutants in which chromosome 3 is absent. (b) Colony-forming units (CFU/ml) of wild-type *Bso* MF6, 34F7, and 27D6 grown alone or in co-culture with Chlamydomonas (+C) for two days. Boxes show median and interquartile range; points are individual colony counts (*n* = 8–12 per group). Significance is Welch’s t-test on log₁₀(CFU) with Holm correction across the five comparisons shown: co-culture reduces the CFU of both mutants (34F7, *p* = 2.8 × 10⁻⁵; 27D6, *p* = 1.5 × 10⁻⁴) but not of wild-type (*p* = 1.00), and neither mutant differs from wild-type when grown alone (34F7, *p* = 1.00; 27D6, *p* = 0.74). (c) Principal-component analysis of the *Bso* MF6 proteome for wild-type, 27D6, and 34F7, grown alone (open symbols) or in co-culture with Chlamydomonas (filled symbols), at day 2 (triangles) and day 3 (circles). Each point is one biological replicate. (d) Abundance of candidates across genotypes and conditions. Rows, wild-type *Bso* MF6, 27D6 and 34F7 grown alone (−) or in co-culture with Chlamydomonas (+); columns, individual proteins with locus tags in parentheses. Color indicates mean log₂ normalized LFQ intensity across four biological replicates; light grey, not detected; hatched, protein encoded on chromosome 3, which is absent from these mutants. Bolded names are considered promising targets. Green marks in the bottom row indicate proteins that are ON or up in the supernatant of MF6 co-cultured with Chlamydomonas (see f). (e) Principal-component analysis of the culture supernatant (Sup) proteome of wild-type *Bso* MF6 (green), 27D6 (orange) and 34F7 (blue) grown alone (open symbols) or in co-culture with Chlamydomonas (filled symbols) at day 2. Each point is one biological replicate. (f) Volcano plot of differential protein abundance in the culture supernatant of wild-type *Bso* MF6 in co-culture with Chlamydomonas versus monoculture at day 2. Dashed lines mark |log₂FC| = 1 and adjusted p = 0.05. Proteins are shaped and colored by subcellular localization predicted with DeepLocPro: extracellular (green crosses, n = 18), outer membrane (orange triangles, n = 25) and periplasmic (blue squares, n = 23); all other proteins are shown in grey. Red circles mark proteins that are up (log₂FC > 1, adjusted p < 0.05) or ON in MF6 only, and not in 27D6 or 34F7. The strip above the plot shows proteins detected in only one condition (“ON in MF6 Sup” or “ON in MF6 +C Sup”), for which fold change cannot be calculated. Position along the strip indicates the percentile of the group the protein is missing from (50 at the centre, 100 at each edge). Selected proteins are labeled with their locus tags in parentheses; bold names are promising candidates from (c). See also Extended Data Figs. 3 and 4 and Supplementary Tables S12-S16.

Additional disrupted genes (Extended Data Fig. 3f) span amino acid biosynthesis, carbon catabolite regulation, protein folding, RNA turnover and a hypothetical protein, mirroring the enrichment for catabolism observed in the proteomics data (Fig. 2c). Of these six, only two had also lost chromosome 3. Although several of these genes act in central metabolism, all six formed visible colonies on Chlamydomonas lawns, indicating that their loss of inhibition does not reflect a general growth defect (Extended Data Fig. 3e).

We compared the proteomes of T6SS-related transposon mutants 34F7 and 27D6 to that of the wild-type, in monoculture and in co-culture with Chlamydomonas, focusing on toxin- and secretion-related proteins. Two days after co-inoculation, while the wild-type maintained a similar colony forming unit (CFU) count in the presence and absence of Chlamydomonas, the mutants 34F7 and 27D6 exhibited a significant reduction in CFU count, only under co-culture with Chlamydomonas (Fig. 3b), indicating loss of a competitive advantage. PCA demonstrates that while the genotype effect is apparent in PC1, co-culture with Chlamydomonas altered *Bso* MF6 proteome along PC2, an effect that is weakened in the mutants in day 2 and abolished by day 3. The proteomes of both mutants shifted away from wild-type mostly when co-cultured with Chlamydomonas (Fig. 3c).

Hierarchical clustering resolved eight co-abundant clusters (1690 proteins out of 4023 in total, Extended Data Fig. 4a and Supplementary Table S9). The enrichment patterns of clusters 2 and 5, that were induced in wild-type co-cultures and not in the mutants, indicate a sensing, uptake and proteostasis response to Chlamydomonas that the mutants fail to mount. In pairwise comparisons (Extended Data Fig. 4b, c), 27D6 showed a relatively minor proteomic shift in monoculture compared with wild-type (107 of 3,246 quantified proteins differing at log2FC > 1, *q* < 0.05) whereas 34F7 shifted more substantially (339 of 3,131 quantified proteins at log2FC > 1, *q* < 0.05).

We identified 34 proteins (Fig. 3d, Supplementary Table S14) that: (i) contain a toxin domain^26^, a T6SS effector, or are predicted to be extracellular^27^; and (ii) were enriched in wild-type (and not in the mutants) in the presence of Chlamydomonas (Fig. 2c). To specifically target extracellular proteins, we also analyzed the proteomes of the culture supernatants (secretome). PCA of these samples mirrors that of cell pellets (Fig. 3e), but with stronger clustering, indicating that secreted proteins are more affected by the interaction with Chlamydomonas. Interestingly, most of the secretome proteins that are upregulated in *Bso* MF6 in the presence of Chlamydomonas are gone in both mutants (Fig. 3f, Extended Data Fig. 4d, Supplementary Tables S15 and S16). Seven of these are among the prioritized list of 34 proteins (Fig. 3d, Supplementary Table S14). The T6SS effector was not among these, possibly because it is directly injected into the target. Two secreted metallopeptidases are in particular interest due to domain architecture and genomic context (Fig. 3f, Supplementary Table S14). Taken together, genetic and proteomic evidence indicate a multi-pronged extracellular attack mounted by *Bso* MF6 against Chlamydomonas that is attenuated in both mutants. This attack involves deployment of the Tad system, a T6SS effector and two secreted metallopeptidases.

### Commonalities and differences in *Bso* MF6 inhibition towards plants and algae

We next asked whether the same mechanism underlies inhibition of Chlamydomonas and Arabidopsis. We inoculated the 14 non-inhibitory mutants onto one-week-old seedlings and assessed inhibition of both root and shoot growth. Eight of the 14 mutants tested had at least partially lost root and shoot growth inhibition capacity (Fig. 4a-c; Supplementary Tables S17 and S18), with five exhibiting complete loss of shoot growth inhibition. In the case of 36A7 this loss of inhibition was accompanied by loss of plant colonization capacity, but not so in the other four mutants (Fig. 4d). Taken together, the causal identification of 11 genes involved in inhibition of Chlamydomonas, and the overlap with inhibition towards Arabidopsis, show that *Bso* MF6 inhibits both Chlamydomonas and Arabidopsis at close proximity, employing multiple modes of action, some of which overlap between unicellular algae and vascular plants.

**Figure 4.**
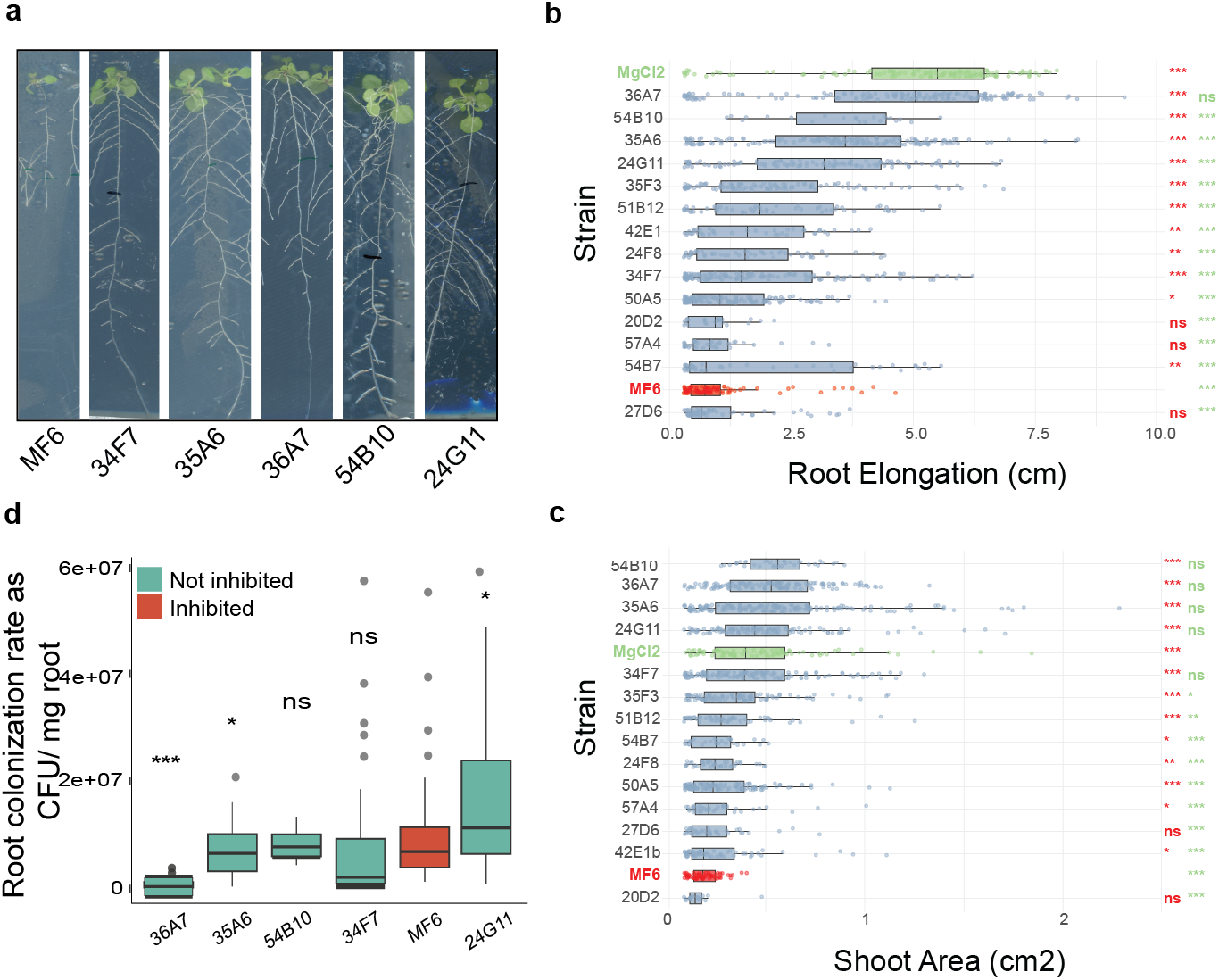
Transposon mutagenesis identifies *Bso* MF6 loci required for inhibition of Chlamydomonas and Arabidopsis. (a) Representative Arabidopsis seedlings 12 days after transfer onto plates inoculated with wild-type *Bso* MF6 or the indicated transposon mutants. (b and c) Primary root length (b) and rosette area (c) of Arabidopsis seedlings 12 days after transfer to plates inoculated with wild-type *Bso* MF6 (red), individual transposon insertion mutants (blue), or MgCl₂ buffer alone (green). Strains are ordered by median value. Boxes show the median and interquartile range (IQR), whiskers extend to the most extreme values within 1.5 × IQR, and points represent individual seedlings. Data are pooled from five independent experiments. Each mutant was compared to wild-type MF6 by Welch’s *t*-test with Benjamini–Hochberg correction; asterisks denote adjusted *p* < 0.05. *n* = 22–196 seedlings per group in (b) and 35–186 in (c). (d) Root colonization of Arabidopsis by wild-type *Bso* MF6 and five Tn5 insertion mutants, expressed as colony-forming units per mg root. Strains are colored by whether they still inhibit root growth (red) or have lost inhibition (teal). Boxes show the median and interquartile range, whiskers extend to 1.5× the interquartile range, and points are outliers (*Bso* MF6, *n* = 52; 24G11, *n* = 58; 34F7, *n* = 38; 35A6, *n* = 73; 36A7, *n* = 74; 54B10, *n* = 4). Each mutant was compared with wild-type *Bso* MF6 by two-sided Student’s *t*-test (\**p* < 0.05, \*\**p* < 0.01, \*\*\**p* < 0.001; *ns*, not significant). See also Supplementary Tables S17-S18.

### Chlamydomonas undergoes regulated cell death upon infection with *Bso* MF6

We next asked what physiological response Chlamydomonas mounts upon infection. We measured gene expression in a mid-log culture (0, 1, 2, 3 and 4 h post-infection) and observed a progressive shift in the transcriptional profile (Fig. 5a-c; Supplementary Tables S19, S20). Temporally transcribed genes fell into 12 clusters (six up- and six downregulated) differing in peak response time and general trend (Fig. 5a, b). Two clusters were sharply and transiently induced at one hour and returned to or below baseline thereafter (clusters 1 and 12; Fig. 5a, b), which we consider the acute response. These clusters were dominated by microtubule- and flagellum-related terms alongside calcium ion binding (Extended Data Fig. 5a, b, Supplementary Tables S21 and S22). The composition, magnitude and kinetics of the response resemble flagellar regeneration following mechanical or acid-shock deflagellation^28–32^. Since calcium influx is the established trigger for flagellar abscission^33^, we interpret the acute response as rapid loss of flagella upon infection, followed by an attempt at regeneration.

**Figure 5.**
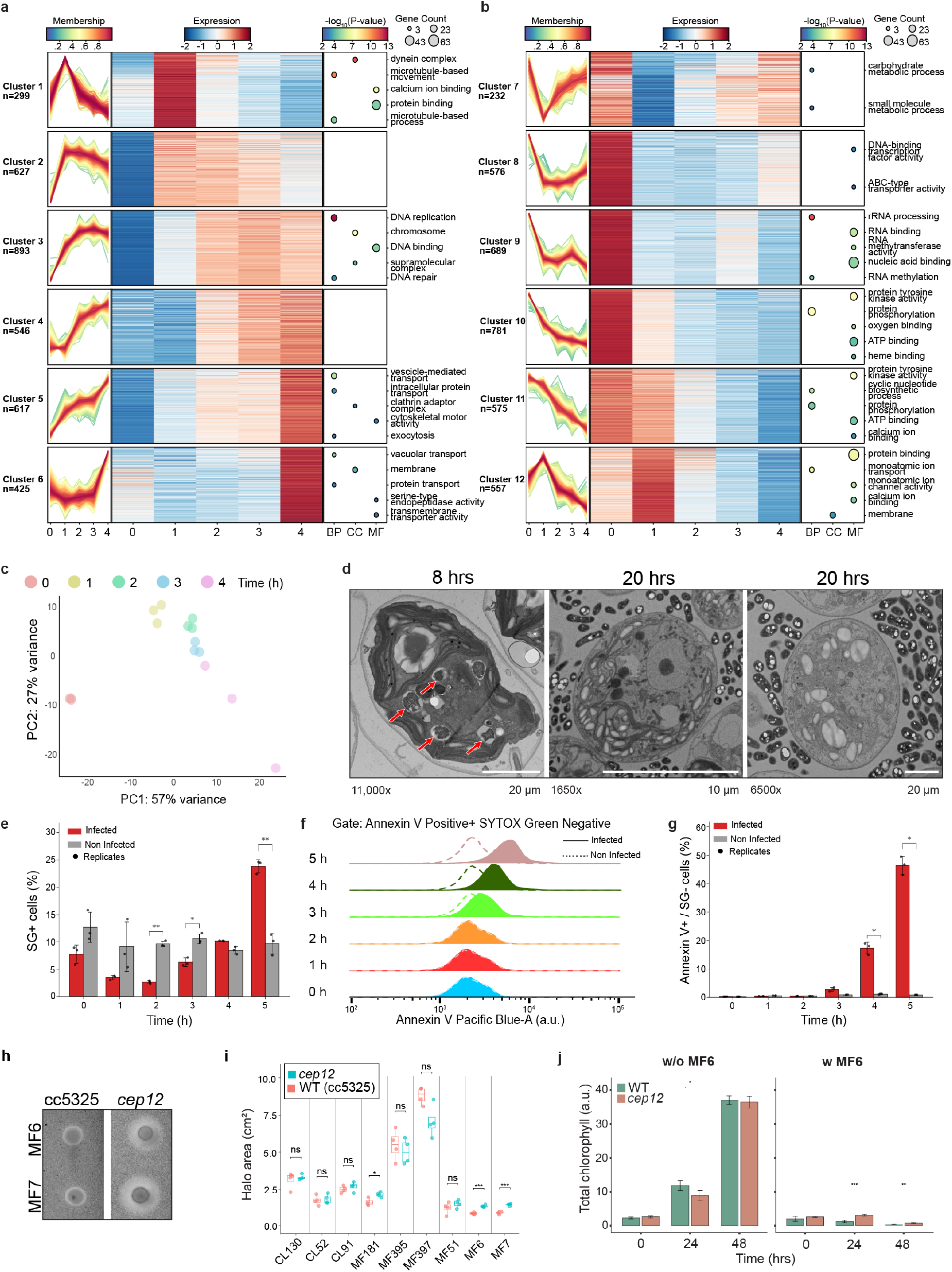
Infection triggers a regulated, non-lytic cell death. (a and b) Temporal clustering and functional enrichment of genes upregulated (a, clusters 1–6) and downregulated (b, clusters 7–12) during infection of Chlamydomonas with *Bso* MF6. For each cluster: left, expression trajectories of individual genes across 0–4 h post-infection, colored by fuzzy c-means membership score; center, heatmap of standardized expression per gene at each timepoint; right, enriched Gene Ontology terms (BP, biological process; CC, cellular component; MF, molecular function), with point size indicating gene count and color the −log₁₀ *p* value (topGO weight01 algorithm, Fisher’s exact test). Cluster sizes are indicated. (b) Principal-component analysis of Chlamydomonas transcriptomes across infection, based on the 1,000 most variable genes after rlog transformation. Points are colored by time post-infection; each point represents one biological replicate. (d) Transmission electron micrographs of Chlamydomonas cells 8 h (left) and 20 h (center, right) after inoculation with *Bso* MF6. At 8 h, cells show convoluted internal membranes and electron-lucent vesicular and vacuolar profiles (arrows). At 20 h, the cytoplasm is largely occupied by large electron-lucent vacuoles with loss of recognizable internal organization, while bacteria remain extracellular; the cell boundary remains continuous. accelerating voltage 120 kV. Magnification (11,000x, 1650x, 6500x) and scale bars (20 µm, 10 µm, 20 µm) from left to right. (e) Percentage of SYTOX Green-positive (SG⁺) cells, indicating loss of plasma membrane integrity, in infected (red) and uninfected (grey) Chlamydomonas cultures over 5 h. Bars show the mean, error bars [SD], and points individual biological replicates (*n* = 3). (f) Annexin V–Pacific Blue fluorescence distributions of SYTOX Green-negative cells at each timepoint, in infected (solid) and uninfected (dashed) cultures. (g) Percentage of Annexin V-positive/SYTOX Green-negative cells, indicating phosphatidylserine exposure in cells retaining membrane integrity, in infected (red) and uninfected (grey) cultures. Bars show the mean, error bars [SD], and points individual biological replicates (*n* = 3). In (e) and (g), infected and uninfected cultures were compared at each timepoint by Welch’s *t*-test with Holm–Šídák correction across timepoints (\**p* < 0.05, \*\**p* < 0.01, \*\*\**p* < 0.001). (h) Representative halo assays of the parental strain CC-5325 and the *cep12* insertional mutant, spotted with *Bso* MF6 (top) or *Bso* MF7 (bottom). Cut from the full panel in Figure S4F. (i) Halo area on lawns of CC-5325 and *cep12* challenged with nine bacterial isolates. [unpaired Student’s *t*-test, adjusted *p* values (Benjamini–Hochberg)]. (j) Total chlorophyll (arbitrary units) over 48 h, alone (w/o *Bso* MF6) or in co-culture with *Bso* MF6 (w MF6), for wild-type and *cep12*. [ANOVA followed by Tukey, *n* = 4]. See also Extended Data Fig. 5 and Supplementary Tables S19-S25.

In contrast to clusters 1 and 12, clusters 2-11 changed progressively and split into two coherent programs. The induced clusters are enriched for vesicle-mediated transport, exocytosis, vacuolar transport and serine-type endopeptidase activity, with induction of ESCRT and retromer subunits and an intensifying ER signature, alongside enrichment for DNA replication and repair (Fig. 5a and Extended Data Fig. 5a). The repressed clusters exhibit carbohydrate and small-molecule metabolism drop, followed by loss of rRNA processing and ribosome biogenesis, consistent with the growth arrest observed in co-culture. Protein phosphorylation and ATP binding decrease in parallel (Fig. 5b and Extended Data Fig. 5a).

Against this background, the vacuolar processing enzyme VPE1 was induced in the progressive phase (cluster 6; Extended Data Fig. 5c, d). Vacuolar processing enzymes are established mediators of plant PCD during pathogen attack^34,35^, making VPE1 a direct death-associated transcript in the dataset. Additional genes implicated in cell death in Chlamydomonas, plants and budding yeast were also upregulated in this phase (Extended Data Fig. 5c, d; Supplementary Tables S23 and S24)^36–40^.

In TEM images, 8 h post-infection, cells displayed convoluted internal membranes and electron-lucent vesicular and vacuolar profiles (arrows, Fig. 5d), and by 20 h the cytoplasm was occupied by large electron-lucent vacuoles with loss of recognizable internal organization, while bacteria had proliferated extensively in the surrounding medium but remained extracellular. Notably, degraded cells retained a continuous cell boundary and no signs of lysis.

Both vacuolar signature in the transcriptome and vesicular profiles seen by microscopy could reflect induction of autophagy as a survival mechanism. However, autophagy-related genes exhibited a mixed and non-canonical transcriptional pattern (Extended Data Fig. 5e; Supplementary Table S25). Notably, the ATG8 conjugation components required for canonical autophagosome formation^41^, ATG3 and ATG7, were both significantly repressed, supporting an alternative degradative program.

We thus hypothesized that induction of VPE1, together with the non-lytic ultrastructure we observed in TEM, represent a regulated cell death program. To test this, we followed phosphatidylserine exposure and plasma membrane integrity by flow cytometry (Fig. 5e-g). Loss of membrane integrity in infected cells was not detectable until 5 h post-infection, when SYTOX Green-positive cells reached 24% versus 10% in uninfected cultures (Fig. 5e). By contrast, Annexin V binding increased markedly earlier, with the Annexin V-positive/SYTOX Green-negative population rising to 18% at 4 h and 46% at 5 h (Fig. 5f, g).

Phosphatidylserine exposure therefore precedes permeabilization, a pattern consistent with a regulated degradative death, in agreement with the intact cell boundaries seen by electron microscopy (Fig. 5d). Together, these transcriptional, ultrastructural, and biochemical signatures indicate that infection triggers two sequential responses: rapid deflagellation followed by an ordered, regulated death.

We next asked whether the conserved algal cell death genes^36^ were engaged during infection. Among these is the cysteine endopeptidase *CEP12* (Cre12.g498850) (Extended Data Fig. 5d), encoding the Chlamydomonas cathepsin X/Z ortholog, for which cell death function has been experimentally demonstrated^36^. In our experiment, *CEP12* was significantly repressed at every timepoint of infection (cluster 9; Fig. 5b). Because *CEP12* is known to be induced under oxidative stress, and its loss confers resilience to oxidative-stress-induced cell death^36^, its repression here, during a lethal infection, was unexpected, and prompted us to test its contribution genetically. Here, the *cep12* insertional mutant was consistently *more* susceptible to *Bso* MF6 and MF7 than its parental strain (Fig. 5h, i and Extended Data Fig. 5f), with only a minor effect on other isolates (Fig. 5i). Notably, this hyper sensitivity was not demonstrated in liquid co-culture with *Bso* MF6 (Fig. 5j). Cathepsin-X therefore acts as a protective factor specifically in the context of cell death induced by a bacterial infection (Fig. 5a and Extended Data Fig. 5a).

### Forward genetics identifies ZIP3 as a susceptibility factor epistatic to CEP12

Next, to identify algal genes required for inhibition by *Bso* MF6 without prior assumptions, we hunted for resistant mutants on bacterial lawns following UV mutagenesis (Extended Data Fig. 6a). Two independent mutants showed reduced halo formation against *Bso* MF6 (Fig. 6a). In both, resistance segregated 2:2 among progeny of crosses to wild-type, consistent with a single causal genetic lesion: tetrad analysis showed a parental ditype (PD):tetratype (TT):non-parental ditype (NPD) ratio of 8:0:0 for *zip3-1* and 9:0:0 for *zip3-2* (Extended Data Fig. 6b, c). Bulked-segregant analysis (BSA)^42^ mapped the causative region in both independent mutants to the same gene locus (Fig. 6b and Extended Data Fig. 6d). Sequencing identified two distinct mutations in *ZIP3* (Cre03.g189550), encoding a ZIP-family zinc/iron transporter: *zip3-1* carries a missense substitution and *zip3-2* a nonsense mutation (Fig. 6b). Complementation of *zip3-2* with the wild-type gene restored susceptibility (Fig. 6c), establishing *ZIP3* as causal. Across a panel of nine bacterial isolates, *zip3* mutants were significantly more resistant only to *Bso* MF6, MF7 and, to a lesser extent, *Pseudomonas* MF51 (Fig. 6d and Extended Data Fig. 6e). Notably, this resilience was not demonstrated in liquid co-culture (Fig. 6e), indicating that the phenotype has a structural, multicellular component. In the transcriptome *ZIP3* was induced in the progressive phase (cluster 4; Fig. 6f; Supplementary Table S26), alongside the vacuolar signatures. This occurred against a broader reorganization of metal homeostasis (Fig. 6f): the zinc-nutrition-responsive plasma-membrane transporters were strongly repressed (ZRT1/2/4/5, IRT2, and ZIP1, clusters 9-11; Fig. 5b), as were the metal tolerance proteins MTP1/3/5, whereas the remaining ZIP transporters were induced (ZIP6, cluster 1; ZIP3, cluster 4; ZIP7, cluster 6; Fig. 5a). Reduced uptake and reduced vacuolar sequestration alongside increased intracellular transport suggests that infection is accompanied by cellular redistribution of zinc/iron. Since *ZIP3* encodes a metal transporter, we asked whether metal availability influences the interaction. Halo formation was strongly increased when omitting Hutner’s trace elements, abolishing the significant effect of *zip3* over WT (Extended Data Fig. 6f, g). We also found that wild-type becomes hypersensitive to infection when iron, but not zinc, is depleted (Extended Data Fig. 6h-j). This effect is mostly abolished in the *zip3* mutants, which maintained resistance under all conditions tested, although iron or zinc omission slightly reduced their resistance phenotype. The ZIP3 complementation line on the other hand, showed hypersensitivity, presumably due to overexpression of *ZIP3* (Extended Data Fig. 6i). Thus, low metal availability exacerbates bacterial inhibition in a *ZIP3*-dependent manner.

**Figure 6.**
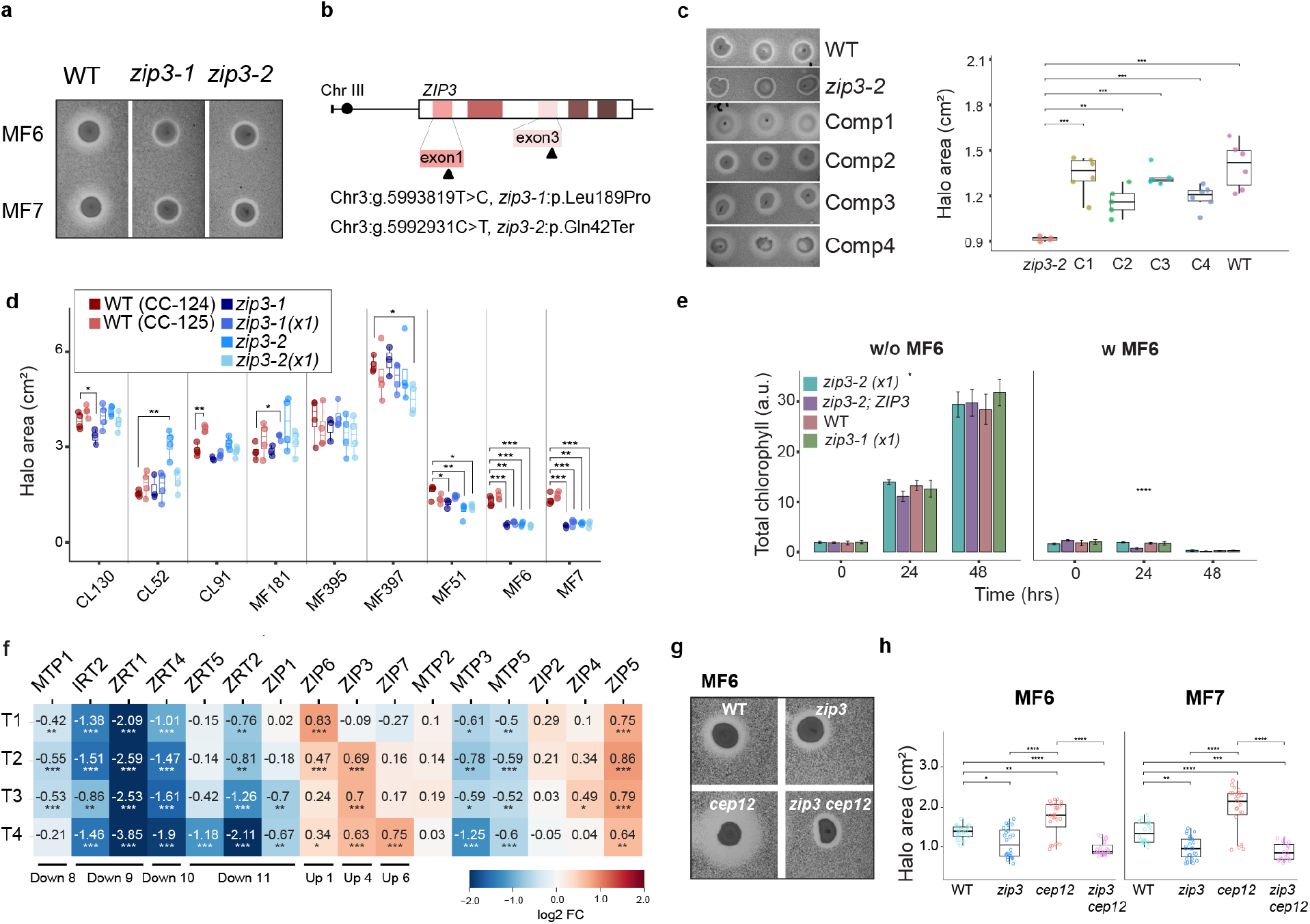
ZIP3 is required for inhibition by *Burkholderia sola* MF6 and MF7 and is epistatic to CEP12. (a) Representative halo assays of wild-type Chlamydomonas and the two UV-derived resistant mutants *zip3-1* and *zip3-2*, spotted with *Bso* MF6 (top) or *Bso* MF7 (bottom). Cut from the full panel in Figure S5F. (b) Position of the causal lesions within *ZIP3* (Cre03.g189550) on chromosome 3. Boxes denote exons; arrowheads mark the *zip3-1* missense substitution in exon 1 and the *zip3-2* nonsense mutation in exon 3, with genomic and protein-level annotations shown below. (c) Complementation of zip3-2 restores susceptibility to *Bso* MF6. (Left) Representative halos formed by *Bso* MF6 on lawns of wild-type, *zip3-2*, and four independently complemented transformants (Comp1–4). (Right) Quantification of halo area. Boxes show median and interquartile range, whiskers extend to the most extreme value within 1.5 × IQR, and points show individual replicates (*n* = 4, 6, 5, 5, 6, 6 for wild-type, zip3-2, and Comp1–Comp4, respectively). Halo areas were compared with zip3-2 by unpaired two-tailed Student’s *t*-test; adjusted *p* values (Benjamini–Hochberg) are shown. See also Online Methods. (d) Halo area on lawns of two wild-type backgrounds (CC-124, CC-125), the two *zip3* mutants, and their single-backcross progeny (x1), challenged with nine bacterial isolates. Boxes show median and interquartile range; points are individual replicates. [unpaired Student’s *t*-test, adjusted *p* values (Benjamini–Hochberg)]. (e) Total chlorophyll (arbitrary units) over 48 h in liquid culture, alone (w/o MF6) or in co-culture with *Bso* MF6 (w MF6), for *zip3-2*(x1), the *zip3-2*;ZIP3 complemented line, wild-type, and *zip3-1*(x1) [ANOVA followed by Tukey, n=4]. (f) Expression of metal-transporter genes across infection (T1–T4 vs. T0). Numbers are log₂ fold change; color scale as indicated. Asterisks denote adjusted significance from DESeq2 (\**p* < 0.05, \*\**p* < 0.01, \*\*\**p* < 0.001). Bars beneath the heatmap indicate the temporal cluster to which each gene was assigned (Figure 4A and 4B). (g) Representative halos formed by *Bso* MF6 on lawns of wild-type, *zip3*, *cep12* and the *zip3 cep12* double mutant. (h) Halo area for the four genotypes in (h) challenged with *Bso* MF6 (left) and *Bso* MF7 (right). [one-way ANOVA followed by Tukey’s HSD] See also Extended Data Fig. 6 and Supplementary Table S26.

ZIP3’s closest homolog in Arabidopsis is ZTP29 (AT3G20870, Extended Data Fig. 6k, l). However, *ztp29* mutants in Arabidopsis did not show reduced susceptibility to *Bso* MF6 compared to wild-type (Extended Data Fig. 6m). This may reflect redundancy within the Arabidopsis ZIP family. Finally, to ask how *ZIP3* and *CEP12* interact, we constructed a *zip3 cep12* double mutant. The double mutant was indistinguishable from *zip3* alone and significantly more resistant than either *cep12* or wild-type, for both *Bso* MF6 and MF7 (Fig. 6g, h). Notably, halo areas were comparable between the walled wild-type strain (CC-124) and the cell-wall-deficient strain CC-5325 (Fig. 5i and 5d), indicating that strain background and wall status do not affect susceptibility. *ZIP3* is therefore required for the increased susceptibility of the *cep12* mutant. Together, loss of ZIP3 attenuates infection-induced inhibition, whereas loss of cathepsin-X accelerates it, consistent with the repression of CEP12 observed during infection.

## Discussion

Our results establish that a unicellular green algae can serve as a tractable proxy for quantifying plant antagonistic potential among bacteria at scale. The Chlamydomonas phycosphere was recently proposed as a model for the plant rhizosphere on the basis of community composition^16,43^, and we show that this correspondence extends to function. In our screen nearly all Chlamydomonas-inhibiting bacteria also inhibited Arabidopsis. We also show that algal Inhibition can be used to enrich plant antagonists from uncharacterized communities. Using Chlamydomonas, thousands of interactions can be assayed in a single experiment. Our own bacterial mutant screen assayed 5,023 genotypes against the algae. Modern co-culturing platforms can increase this number to hundreds of thousands, orders of magnitude beyond what is feasible in Arabidopsis^44^.

The correlation we observed, though, is asymmetric. More bacteria inhibited Arabidopsis than Chlamydomonas. This may indicate that a unicellular algae is intrinsically more resilient than its multicellular counterpart, or that it lacks the tissues and organs through which antagonists act; it may also reflect our use of an Arabidopsis-derived collection, which is enriched for strains adapted to that host. Either way, Chlamydomonas functions as a high-specificity but incomplete-sensitivity filter, and screens built on it will recover a conserved subset of plant antagonists while missing those that require a root, vasculature, or immune components the algae does not possess.

We also expect antagonism to show graded host specificity, as acute pathogens do^45^; our target-range experiments span a wide range, from *Paenibacillus* MF181, which inhibited bacteria and yeast as well, to *Burkholderia* and *Pseudomonas* strains whose activity was confined to the photosynthetic targets tested. Future infection assays across multiple algal and plant hosts would resolve specificity from generality within photosynthetic lineages.

Features that render both Chlorophytes and land plants susceptible to the same bacteria were most plausibly present in their last common ancestor >470 mya^46^. This contrasts with mutualism, which in several cases appears to be recently derived from pathogenic ancestry^47^. An exploitative lifestyle may thus be the ancestral strategy, with mutualism arising repeatedly and recently from it.

*Burkholderia* are found at the deepest core of the microbiota of terrestrial photosynthetic organisms, including both plants and algae, more so than any other bacterial genus.^48^ We identified the systems that *Bso* MF6 deploys against Chlamydomonas, several of which are shared with its attack on Arabidopsis. The cross-host overlap therefore manifests at both the ecological and the molecular level. The bacterium derives two benefits. It wins direct competition and it gains access to the nutrients released by the dying algae, evident by the coordinate induction of chemotaxis, transport, and catabolic pathways, a sensing-and-foraging response.

The bacterial attack on Chlamydomonas relies on multiple loci. The *Bso* MF6 genome encodes 27 secretion systems spanning nine types, indicating that protein export is central to its survival strategy. 14 non-inhibitory mutants span secretion, central metabolism, protein folding and RNA turnover, and the proteomic comparison of 27D6 and 34F7 with wild-type shows reduced expression of multiple candidate toxins (Fig. 3c). In our transposon mutant screen, chromosome-3 loss co-occurs with T6SS-linked insertion. This therefore implicates T6SS-mediated delivery and chromosome-3 gene content jointly, and cannot establish that either is sufficient alone. This also raises the intriguing possibility that the two are causally linked. Evidence is accumulating for the importance of T6SS in plant-microbe interactions, although direct evidence is still lacking, in our work and in the field generally^49^. Our electron micrographs place bacteria against a locally distorted algal surface with apparent introduction of bacterial material, which is consistent with contact-dependent delivery.

To our knowledge this is the first implication of the Tad system in bacterial antagonism toward plants or algae. Tad is a type IVc pilus system architecturally related to the type II secretion machinery (T2SS, itself also present in the *Bso* MF6 genome and expressed here). T2SS was previously implicated as critical in other cryptic antagonists: *Xanthomonas* strains Leaf131 and Leaf148 are commensals on wild-type Arabidopsis but degrade plant tissue in immunocompromised plants (when other bacteria are absent) via a T2SS-dependent export of cell-wall-degrading enzymes^13^. *Dyella japonica* MF79 (which does not inhibit Chlamydomonas) employs T2SS-secreted proteins to attenuate Arabidopsis immunity^3^. All of these strains were isolated from asymptomatic Arabidopsis roots and leaves^50,51^, and can thus be considered cryptic antagonists.

Interestingly, while *Xanthomonas* virulence relies on plant cell wall-polymer enzymes, *Bso* MF6 has no equivalent in its genome. Instead, *Bso* MF6 induces proteolytic and lipolytic enzymes, which are more suitable for the Chlamydomonas cell wall, that is glycoprotein rich and containing neither cellulose nor pectin^52^. This suggests why *Bso* MF6 antagonism transfers across the green linage. The secreted proteases and lipases in our list of 34 candidates (Fig. 3d) are not as specific to host wall chemistry. We would therefore predict that strains relying on secreted wall-polymer enzymes inhibit land plants without inhibiting Chlamydomonas. In fact, most of the prioritized candidate proteins in *Bso* MF6 (Fig. 3d) are degradative enzymes rather than recognizable toxins. Only the two RHS repeat proteins and the T6SS effector carry canonical toxin architecture. Since these proteins are reduced or absent in 27D6 and 34F7 mutants that lose inhibition, it argues that they are not merely saprophytic. Seven of these shortlisted proteins were also detected in the cell-free supernatant of *Bso* MF6 grown with Chlamydomonas (Fig. 3d), placing them where they would encounter the algae, including two metallopeptidases.

In the host, we interpreted the acute response to infection as loss of flagella on contact followed by attempted regeneration. Deflagellation has precedent as a bacterial strategy in Chlamydomonas: *Pseudomonas protegens* secretes a cyclic lipopeptide that disrupts cytosolic Ca²⁺ homeostasis and immobilizes Chlamydomonas^53^, acting through transient receptor potential-type Ca²⁺ channels^54,55^. The mechanism is different here, since *Bso* MF6 inhibition is largely contact-dependent and its genome encodes no comparable lipopeptide cluster. Whether deflagellation is the *Bso* MF6 strategy to immobilize the host, a host response that discards adherent bacteria along with the flagella, is unresolved.

Our finding that CEP12, a cathepsin X-type cysteine endopeptidase, confers resistance to *Bso* MF6 inhibition (Fig. 5h, i) adds a dimension to our understanding of protease-mediated cell death in photosynthetic organisms. Cathepsin B participates in hypersensitive-response execution in plants, and vacuolar processing enzymes are established mediators of plant PCD during pathogen attack^34,35^. Cathepsin X-type proteases, by contrast, have been characterized primarily in animal systems, in contexts such as inflammation and neurodegeneration^56^. In Chlamydomonas, cathepsin X was identified as a pro-death protein as it is induced under oxidative stress, and its loss confers resistance to H₂O₂-induced cell death^36^. In contrast to this, in our data CEP12 was repressed by infection, and the *cep12* mutant was consistently more susceptible to *Bso* MF6 and MF7 (Fig. 5i and Extended Data Fig. 5f). The dual function proposed for cathepsin X may resolve this: alongside its pro-death role under oxidative stress, *cep12* mutants show impaired stationary-phase carrying capacity and increased sensitivity to several other stressors, suggesting a pro-survival role in protein turnover and recycling^36^. Our data provide direct evidence for that second function during a biotic interaction. To our knowledge this is the first evidence implicating a cathepsin X-type protease in the response of a photosynthetic organism to bacterial attack, raising the possibility that the family plays a broader and previously unappreciated role in green-lineage immunity.

The independent isolation of two *zip3* alleles in a forward genetic screen is a strong evidence for causality, but raises the question for why no other genes in the pathway were recovered. The most likely explanation is that UV generates point mutations, so only lesions that inactivate a protein without being lethal are recovered.

The hallmarks of a regulated cell death exhibited by Chlamydomonas upon infection, together with *zip3* mutant phenotypes being restricted to solid media, point to a multicellular response of Chlamydomonas cultures to infection^57^. An attempt of the Chlamydomonas culture to contain the infection is reminiscent of a vascular plant’s hypersensitive response to infection, also involving calcium bursts^58^, and it is enticing to speculate of a deep evolutionary link between the two.

We therefore propose that *Bso* MF6 engages a ZIP3-dependent, zinc-associated death pathway acting in the endomembrane compartment, and that cathepsin X restrains this pathway, such that its loss accelerates inhibition while loss of ZIP3 blocks it.

Omitting trace elements increased halo formation and abolished the *zip3* advantage, iron depletion rendered wild-type hypersensitive, supporting our observation that the pathogenicity is dependent on secreted metallopeptidases. Iron-availability plays a critical role in host-microbe interactions^59,60^ and our data suggests that this theme extends also to cases where the host is unicellular.

### Limitations of the study

Our bacterial genetics are correlative at the level of individual genes. Chromosome-3 loss and T6SS-linked insertion co-occur in every prioritized mutant, so we cannot determine whether either lesion is sufficient alone. Our candidate toxins were prioritized on domain architecture, genomic context and differential abundance; but none has been shown directly to inhibit. Finally, despite the strong overlap, the algae cannot fully substitute the plant. Prediction from Chlamydomonas to Arabidopsis held in our hands, but the converse does not hold, as Chlamydomonas lacks tissue organization, a root, and much of the immune repertoire of a land plant. Antagonists acting through those features will be invisible to this platform. Our Arabidopsis assays were also conducted on agar under gnotobiotic conditions with single strains, which removes the community context that ordinarily buffers cryptic antagonism in soil; the effect of the cryptic antagonists studied here, is mostly, but not completely abolished in the presence of a complex synthetic community^4^. The magnitude of these effects within a natural microbiome remains to be established.

## Online Methods

### Strains maintenance

Chlamydomonas strains CC-4402 (mt+ gamete) and CC-4403 (mt− gamete), CC5325 (mt− gamete) were maintained on Tris Acetate Phosphate (TAP) plates in 21° and under continuous light (50 µE)^61^.

The bacterial strains were derived from a collection generated at the Dangl lab (UNC-CH, NC, USA)^4,62^. Nearly all strains were isolated from surface-sterilized *Arabidopsis thaliana* roots grown in two soils from North Carolina (USA) and are genome-sequenced^51^. Bacteria were stored as 25% glycerol stocks at −80°C. For each experiment, bacterial stocks were streaked onto Lysogeny broth (LB) agar plates and grown at 28°C in the dark for 1-5 days, depending on the strain’s growth rate. Once colonies were observed, plates were stored at 4°C for up to one month and used for subsequent experiments. Prior to each experiment, a single colony was picked and grown overnight in 2 ml LB at 28°C, 168 rpm.

### Halo assay for assessing antagonistic activity via bacterial-Chlamydomonas co-inoculation on soft agar

Twelve ml of TAP soft agar (0.5% agar) were heated to 38°C, and an overnight culture of Chlamydomonas in TAP was added to a final optical density (OD) of 0.1 (10⁶ cells/mL). The soft agar was poured over TAP plates (1.5% agar) and allowed to solidify. Bacterial strains were grown overnight in liquid LB, diluted to OD₆₀₀ = 1, and 4 µl drops were spotted onto the Chlamydomonas soft agar plates (20 bacterial strains per plate). Plates were incubated under continuous light (50 µE) at 21°C (Figure S1B) and scanned on days 3-5 using both a document scanner (Epson Perfection V850 Pro) and a gel imager (Fusion FX, Vilber Lourmat). Inhibition was quantified as the total halo area minus the area of the bacterial colony, using ImageJ/FIJI software.

### Isolation of antagonistic bacteria from environmental soil and root samples

To identify antagonistic strains directly from the environment, we collected soil and root samples from pine forest soil (31°46’21.7“N, 35°11’53.5”E) using sterilized tools into sterile paper bags and processed them immediately in the lab.

For soil processing, 5 ml of each soil sample was resuspended in 30 ml of 10 mM MgCl₂ in a 50 ml Falcon tube, vortexed vigorously for 30 s, and passed through a 100 µm cell strainer to remove large particles, yielding a final volume of 25 ml. Roots were separated from the soil, shaken manually to remove residual soil particles, and washed in 30 ml of 10 mM MgCl₂ by vortexing for 10 s. In a biosafety cabinet, main and lateral root tips were excised and placed into sterile 1.5 ml Eppendorf tubes containing three 4 mm glass beads. Samples were homogenized using a bead beater (Mini-Beadbeater™, Biospec Products) at maximum speed for 2 min, and the resulting root homogenate was diluted into 30 ml of 10 mM MgCl₂.

To screen for antagonistic activity, 96-well plates were prepared with 90 µl of 10 mM MgCl₂ per well. Ten µl of each soil or root suspension was added to the first well and serially diluted 10-fold across five steps. Four µl of each dilution was spotted onto Chlamydomonas soft agar plates (OD = 0.1) and incubated until a visible inhibitory phenotype was observed. As an alternative sampling approach, 50 µl of each sample was also spread directly onto Chlamydomonas lawns.

To confirm the presence of viable bacteria in each sample, 50 µl was plated onto LB agar and incubated at 28°C overnight. When a halo phenotype was observed on Chlamydomonas plates, bacterial colonies within the halo were isolated onto LB agar and incubated for an additional 1–3 days at 28°C; a secondary round of isolation was performed to ensure colony purity. Purified colonies were grown for 1-3 days in liquid LB and stored at −80°C as 25% glycerol stocks. Each isolate was taxonomically identified by sequencing the V3-V5 region of the 16S rRNA gene using primers 338F (5′-ACTCCTACGGGAGGCWGCAG-3′) and 907R (5′-CCGTCAATTCCTTTRAGTTT-3′).

Antagonistic activity of each isolate against Chlamydomonas was confirmed using the halo assay described above. All plates were scanned using a document scanner (Epson Perfection V850 Pro).

### Halo assay using *Escherichia coli* [strain YL151], *Bacillus subtilis* [strain TY79], and *Saccharomyces cerevisiae* [strain BY4741] as target strains

For the soft agar target strain plates, *E. coli* and *B. subtilis* were grown overnight in 2 ml liquid LB at 28°C, and *S. cerevisiae* was grown overnight in 2 ml Yeast Peptone Dextrose (YPD) medium at 28°C. To prepare the 15 ml soft agar lawn (0.5% agar, 45 mm diameter plate), medium was heated to 40°C. The target strain’s overnight culture was diluted to OD = 0.01 (8.0 × 10⁶ cells/mL for *E. coli* and 5.0 × 10⁶ cells/mL for *B. subtilis*) or OD = 0.1 for *S. cerevisiae* (1.5 × 10⁶ cells/mL), then poured over LB agar plates, or YPD agar plates for *S. cerevisiae* (1.5% agar, 45 mm diameter plate). Plates were incubated for 2 h at 37°C. Inhibitory bacteria were grown overnight in LB at 28°C, diluted to OD = 1 in LB, and 4 µl was spotted onto the target strain soft agar plates.

Plates were then incubated at 28°C, and images were taken after 1–2 days using a document scanner (Epson Perfection V850 Pro) and a gel imager (Fusion FX, Vilber Lourmat).

### *Bso* MF6 genome assembly and quality control

Genomic DNA was submitted to Plasmidsaurus (Eugene, OR, USA) for whole-genome sequencing. Amplification-free long-read libraries were prepared using v14 chemistry with sequence-independent tagmentation-based fragmentation, and sequenced on R10.4.1 flow cells under a primer-free protocol. Reads were basecalled with Dorado v4.3 (Oxford Nanopore Technologies, super-accurate model, default Q10 filtering) and quality-filtered with Filtlong v0.2.1^63^, discarding reads shorter than 300 bp and retaining the best 95% by quality. Genome size was estimated with LRGE v0.2.1^64^, and reads were subsampled into multiple sets at Autocycler-determined coverage^65^. Each subsample was assembled with Flye v2.9.6^66^, hifiasm v0.25.0-r726^67^ and Plassembler v1.8.0^68^; low-depth and short contigs were removed, and the assemblies were compressed, clustered, trimmed, resolved and cleaned with Autocycler v0.6.2 to produce a consensus, which was rotated to a canonical start position with dnaapler v1.4.0^69^. Assemblies were polished with Medaka v1.8.0^70^ using the filtered read set.

The resulting draft assembly of the *Bso* MF6 strain was composed of five contigs. Each contig was screened for foreign origin by querying it against the NCBI core_nt database with megablast (BLASTN 2.17.0+)^71^ at default settings. Two contigs, of 33,033 bp and 9,111 bp, had best hits to taxa other than *Burkholderia* and were removed from the final assembly. The 9,111 bp contig matched chromosome XII of *Saccharomyces cerevisiae* strain I-25 (best hit - GenBank: AP045000.1) at 99.22% identity over 100% query coverage. The 33,003 bp contig aligned exclusively to *Micrococcus* spp. (best hit - GenBank: CP083691.1; 99.99% identity, 69% query coverage) and produced no alignment to any *Burkholderia* sequence.

To further support the contaminant nature and to confirm that no products of this contig were detectable *in vivo*, the 33 protein-coding genes annotated by NCBI Prokaryotic Genome Annotation Pipeline (PGAP) v2026-06-18.build8602^72,73^ on the 33,003 bp contig were appended to the target–decoy search database used for the 48-run co-culture proteomics experiment (see Figure 2). No peptide mapping to these sequences was identified in any of the 48 runs at 1% FDR at the PSM, peptide, or protein level.

The final assembly comprises three circular replicons, named chromosome 1–3 in order of decreasing length, totalling 7,755,817 bp with a G+C content of 66.74%. Completeness and contamination were assessed with CheckM2 v1.1.0^74^, resulting in 100% completeness and 0.15% contamination.

### *Bso* MF6 genome annotation

The curated three-replicon assembly was annotated with the PGAP, yielding 7,024 genes: 6,934 CDSs, of which 6,841 encode proteins and 93 are pseudogenes, and 90 RNA genes (six complete copies each of 5S, 16S and 23S rRNA, 68 tRNAs and 4 ncRNAs). Locus tags of the form AC1V0C_##### are used throughout this study and in all supplementary tables.

COG categories and KEGG orthologs were assigned to the predicted proteome with eggNOG-mapper v2.1.12^75^ against the eggNOG v5.0 database^76^. Sequence searches were performed with DIAMOND v2.1.12^77^ in sensitive mode (--sensmode sensitive) with per-query Pfam realignment (--pfam_realign realign). Protein domain and family assignments were obtained with standalone InterProScan v6.0.1^78^ against InterPro database v109.0^79^. Subcellular localization was predicted with DeepLocPro v1.0^27^ in Gram-negative mode and the highest-probability class was taken as the predicted localization.

### Annotation of bacterial protein secretion systems and biosynthetic gene clusters

Protein secretion systems were predicted in *Bso* MF6 and in ten isolated genomes (Table S7) using the TXSScan 1.1.4^80^ model set under MacSyFinder 2.1.6^81^, searching all models simultaneously (--models TXSScan all). Run settings differed between the complete and draft assemblies. For *Bso* MF6, MacSyFinder was run once per replicon with --db-type ordered_replicon and --replicon-topology circular, since all three replicons are circular. For the ten co-isolates, --db-type gembase was used with --replicon-topology linear. Because the co-isolate assemblies are draft-quality, systems whose loci are interrupted by contig boundaries may be missed, so absence of a system in a co-isolate is therefore not evidence of its absence from the genome. A T7SS cluster was defined by whether the bacteria had two WXG100 toxins, after which came an *essC* gene and then a ubiquitin like domain^82^.

Biosynthetic gene clusters were predicted with antiSMASH v8.0.4^83^ using relaxed detection strictness. 23 regions were identified across the three replicons: five on chromosome 1, eleven on chromosome 2 and seven on chromosome 3.

### Construction of the transposon mutant library for *Bso* MF6

The mutant library was constructed following Song and colleagues^84^, with slight modifications, as restriction inhibitors were not required for *Bso* MF6. Briefly, mutagenesis was performed using the EZ-Tn5 Transposase system (LGC Biosearch™ Technologies, UK). Electrocompetent *Bso* MF6 cells were prepared from an overnight starter culture inoculated into fresh LB medium at a ratio of 5 ml culture to 25 ml fresh medium and grown for approximately 3–4 h to mid-log phase (OD ≈ 0.4). Cultures were chilled on ice for 10 min and centrifuged twice at 4,000 × g for 10 min at 4°C. Cell pellets were washed twice by gentle resuspension in 30 ml ice-cold 500 mM sucrose, followed by centrifugation under the same conditions, and finally resuspended in 0.3 ml ice-cold 500 mM sucrose containing 10% glycerol for immediate use in electroporation. For transformation, 1 µl of EZ-Tn5 transposome was added to 200 µl of electrocompetent *Bso* MF6 cells. The mixture was transferred to a pre-chilled 0.2-cm gap electroporation cuvette and pulsed using a Gene Pulser system (2.5 kV, 25 µF, 200 Ω). Immediately after pulsing, 1 ml LB medium was added to the cuvette and cells were gently resuspended by pipetting. The recovered suspension was transferred to tubes containing a total of 4 ml LB and incubated at 28°C with shaking at 180 rpm for 4–6 h. Following recovery, 70 µl of each transformation mixture was spread onto LB agar plates supplemented with trimethoprim (100 ng/ml) and incubated at 28°C for 2–3 days. A total of 5,026 mutants were obtained, grown in LB medium, and stored in 25% glycerol at −80°C for downstream screening.

### Phenotypic screening of the transposon mutant library on Chlamydomonas

To examine mutant phenotypes on Chlamydomonas, the library was grown in 96 well plates with 200 µl LB and incubated overnight at 28 °C. Using the ROTOR+ system (Singer Instruments, UK), bacteria were then applied in three conditions: (i) onto Chlamydomonas soft agar plates at OD = 0.1, or (ii) into liquid Chlamydomonas cultures (TAP medium) at OD = 0.2. (iii) TAP only plates to measure bacterial growth. Plates were incubated under continuous light (50 µE) at 21°C (Figures S3A and S3B) and scanned on days 3–5 using a document scanner (Epson Perfection V850 Pro).

Inhibitory phenotypes were quantified in two ways: (i) manual comparison of soft agar and liquid culture plates to categorize interaction phenotypes, and (ii) quantification of Chlamydomonas pellet size in liquid culture, calculated by tracing the pellet contour and counting the pixels within it. Based on growth and inhibition outcomes across the solid and liquid media assays, we classified interactions into five phenotypes: complete inhibition, partial inhibition (both with visible bacterial growth), no bacterial growth, inhibition in only one assay, and no inhibition.

Mutants selected for secondary screening were those that lost the ability to inhibit Chlamydomonas, showing either no growth or no halo formation. In this second screen, mutants were inoculated onto Chlamydomonas soft agar at a higher volume (10 µl) to exclude effects of slow mutant growth rate. Mutants that still failed to inhibit Chlamydomonas growth despite the increased bacterial density were carried forward to a third and final screen, from which 14 mutants were selected. All 14 mutants showed normal colony growth but no inhibitory activity toward Chlamydomonas.

### Transposon insertion site mapping and chromosome 3 verification

Twelve selected Tn5 mutants were subjected to whole-genome sequencing to map the cassette insertion site and to verify the presence of chromosome 3. Seven mutants (20D2, 24F8, 24G11, 27D6, 42E1, 50A5 and 54B10) were sequenced on Oxford Nanopore (R10.4.1, super-accuracy basecalling) and five (34F7, 35A6, 35F3, 36A7, and 51B12) on Illumina NovaSeq X Plus (151 bp paired-end); 35A6 and 36A7 were each sequenced as two independent libraries, giving 14 libraries in total. To resolve the integration sites, the Tn5 cassette sequence was appended to the *Bso* MF6 assembly. Nanopore reads were aligned using minimap2 v2.30^85^ (-ax map-ont –MD-Y -L --secondary=no), retaining supplementary alignments. Illumina reads were aligned with BWA-MEM 0.7.17^86,87^ (-Y, soft-clipping for supplementary alignments) and processed with samtools fixmate -m, sort and markdup (SAMtools 1.19.2)^88^.

Insertion sites were detected per clone using a custom script implementing two independent detection methods: reads spanning the complete cassette with genomic sequence on both flanks (nanopore only), and cassette junctions, i.e. reads split between a chromosomal sequence and the transposon sequence (both platforms). Evidence from the two detectors was pooled per site and counted per read. Junctions were clustered within 150 bp to absorb indel noise around the breakpoint, and clusters supported by fewer than three (nanopore) or five (Illumina) reads were discarded. Allele fraction was calculated as supporting reads divided by the sum of supporting and reference-spanning reads, where a reference-spanning read carried at least 100 bp (nanopore) or 25 bp (Illumina) of contiguous alignment on each side of the site. Each clone carried a single insertion at an allele fraction of 1.00 (Table S12). Insertion coordinates were determined as the 5’-end of the target site duplication sequence.

Replicon presence was assessed with samtools coverage (SAMtools 1.19.2) applied to the alignments described above, counting only reads with mapping quality ≥ 10. Chromosome 3 was treated as absent when breadth of coverage fell below 1% and present when it exceeded 99%. Eight mutants (20D2, 24G11, 27D6, 34F7, 35F3, 42E1, 50A5 and 51B12) showed breadth of 0.002–0.35% over chromosome 3. Each of the remaining four mutants (24F8, 35A6, 36A7 and 54B10) covered ≥ 99.5% of chromosome 3. Chromosomes 1 and 2 and the transposon contig were covered in full in every clone. Per-clone coverage and breadth values are given in (Table S13).

The integration sites for two additional mutants (54B7 and 57A4) were determined by Sanger sequencing of PCR products amplified with the primer pair 5′-GTGCCTACGAGCTAGCCAAAG-3′ (forward) and 5′-TCATCCCCGAGCGCATC-3′ (reverse), used for both mutants.

### Co-inoculation of Arabidopsis seedlings with single bacterial strains for plant phenotyping

*Arabidopsis thaliana* Col-0 seeds were surface-sterilized in 70% bleach with 0.2% Tween-20, shaking at 1,000 rpm for 15 min in 1.5 ml Eppendorf tubes. The bleach solution was removed by washing five times with autoclaved distilled water, and seeds were resuspended in 1 ml autoclaved distilled water. Seeds were vernalized at 4°C in the dark for 24 h, then germinated on Murashige and Skoog (MS) medium containing 1% sucrose (0.8% agar, 12 × 12 cm square plates) under long-day conditions (16 h/21°C day, 8 h/18°C night) for 7 days.

In parallel, bacteria were grown overnight in 4 ml liquid LB one day prior to transfer. On the day of transfer, bacterial cultures were washed with 10 mM MgCl₂ and diluted to OD = 0.02. One hundred µl of the diluted culture was spread evenly onto fresh 0.5× MS plates (0.8% agar, 12 × 12 cm square plates). Ten 7-day-old seedlings were then transferred from germination plates onto the bacteria-coated plates. Plates were transferred to a short-day regime (8 h/21°C day, 16 h/18°C night) to favor vegetative growth over flowering, and grown for an additional 12 days. Plates were imaged using a document scanner (Epson Perfection V850 Pro), and plant morphology was analyzed using ImageJ^89^. Plants with obvious developmental stunting (e.g. did not develop true leaves) were not measured.

### Measuring bacterial colonization of Chlamydomonas and Arabidopsis roots

Bacterial colony forming units (CFU) were quantified from Arabidopsis roots and from liquid co-culture with Chlamydomonas to evaluate colonization by mutant strains relative to wild-type *Bso* MF6. For root colonization measurements, inoculated roots were collected after 10 days of growth. All roots from each plate were pooled into a tube containing three sterile glass beads and weighed. Two hundred µl of 10 mM MgCl₂ was added to each tube, and samples were homogenized in a bead beater for 30 s. An additional 300 µl of 10 mM MgCl₂ was then added to the homogenized roots, for a final volume of 0.5 ml.

For Chlamydomonas liquid co-culture, 100 µl of the co-culture was collected after 2 days of growth. From both root and co-culture samples, a 1:10 dilution series was prepared in 96-well plates, down to 10⁻⁶. Four µl droplets from each dilution were plated onto LB agar and incubated at 28°C for 1–2 days before colony counting.

### Sample collection and preparation for proteomic analysis

Wild-type *Bso* MF6 was compared with the Tn5 mutants 27D6 and 34F7, each grown alone or in co-culture with Chlamydomonas. Chlamydomonas overnight culture in TAP was diluted in fresh TAP to a final OD of 0.1, and bacterial overnight cultures were washed in TAP and diluted to OD=0.1. Co-cultures were assembled in 50 ml tubes containing 25 ml Chlamydomonas (OD=0.1) and 25 µl bacterial culture (OD=0.1 to reach a final concentration of 0.0001); control tubes containing Chlamydomonas alone were prepared in parallel. Tubes were placed on a rotating stand at ∼50 rpm, 21°C, under continuous light. Samples for proteomics were collected at four timepoints post-inoculation (1, 2, 3, and 4 days). At each timepoint, 2 mL of culture was harvested and placed directly on ice, then centrifuged at 20,000 x g for 2 min. The supernatant was decanted into a new tube, filtered through a 0.22 µm filter, and frozen in liquid nitrogen; the pellet was blotted free of excess liquid and flash-frozen in liquid nitrogen separately. For protein extraction, pellets were resuspended in 100–200 µl lysis buffer (25 mM Tris-HCl, pH 8, supplemented with 4% SDS) and sonicated three times (25% amplitude, 8 s pulses). Lysates were centrifuged at 13,000 × g for 8 min, and the supernatant containing the extracted proteins was transferred to new Eppendorf tubes. Samples were then heated at 95°C for 5 min. Proteins samples were digested and cleaned using S-Trap micro columns (Protifi, LLC, Huntington, NY) according to manufacturer’s protocol. In brief, proteins in 4% SDS/TRIS-HCl 25mM pH8 were reduced by the addition of DTT to a concentration of 10mM and incubated for 30min at room temperature, followed by alkylation in 55 mM iodoacetamide and incubation for 30 minutes at room temperature in the dark. Phosphoric acid was added to a final concentration of 1.2%. Methanol-TRIS buffer (90% MeOH, 10% TRIS 50mM pH7.1) was added to the samples at a ratio of 6:1 (buffer:sample) and loaded onto S-Trap columns by centrifugation at 1,000g for 1 minute until all sample was loaded onto column. Loaded columns were washed twice with 150µl Methanol-TRIS buffer at 4,000g. Sequencing grade modified trypsin (Promega Corp., Madison, WS) was added onto the column (1µg per column) and incubated at 47 C^O^ for 90 minutes. Peptides were eluted from column, acidified and desalted on homemade C18 stage tips^90^.

Peptides were eluted from stagetips and 0.35 µg (determined by Absorbance at 280 nm) from each sample were injected into the mass spectrometer for analysis. Frozen supernatants that were collected through the experiment (1, 2, 3, and 4 days) were lyophilized. Dried conditioned samples were resuspended in 100 μl 8M urea, 10 mM DTT, 25 mM Tris-HCl pH 8.0 and incubated for 30 min at room temperature. Iodoacetamide was added to a final concentration of 55 mM followed by incubation of 30 min in the dark. Eight volumes of 25 mM Tris-HCl pH 8.0 were added to dilute the urea and 0.4 μg/ sample of Trypsin was added and incubated overnight at 37°C with gentle agitation. Next day the peptides were acidified using formic acid to final concentration of 0.38% and desalted on home-made C18 Stage tips. Peptides were eluted using 80% Acetonitrile (ACN), 0.1% Formic acid (FA) then dried and resuspended in 10 μl water containing 0.1% FA from which 1 μl was injected into the mass spectrometer for analysis.

### LC-MS/MS analysis of cell pellets

Analysis was performed at the Stein Family mass spectrometry center in the Silberman Institute of Life Sciences, Hebrew University of Jerusalem. MS analysis of cell pellets was performed using a Q Exactive-HF mass spectrometer (Thermo Fisher Scientific, Waltham, MA USA) coupled on-line to an Ultimate 3000 Dionex (Thermo Fisher Scientific, Waltham, MA USA) UHPLC. Peptides were separated on a gradient (A - 99.9% water, 0.1% Formic acid, B-99.9% acetonitrile 0.1% formic acid) run at a flow rate of 0.30 μl/min from 4%B to 50%B over 112 minutes on a reverse phase 25-cm C18 column (Aurora Ultimate XT 25×75, ionopticks, AU). Gradient continued up to 80% B over 22 minutes followed by thorough washing in 80% B and equilibration in 1% B before the next sample was loaded. Data was acquired using Xcalibur software (Thermo Scientific). Survey scans (350–1,200 m/z, target value 3E6 charges, maximum ion injection time 40 ms) were acquired and followed by higher energy collisional dissociation (HCD) based fragmentation (normalized collision energy 27). A resolution of 120,000 was used for survey scans and up to 15 dynamically chosen most abundant precursor ions were fragmented (isolation window 1.6 m/z). The MS/MS scans were acquired at a resolution of 15,000 (target AGC 1E5, maximum ion injection times 45 ms). Dynamic exclusion was set to 30 sec and peptide preferred profile option selected.

### nanoLC-MS/MS analysis of supernatants

MS analysis was performed using Astral mass spectrometer (Thermo Fisher Scientific, Waltham, MA USA) coupled to Vanquish Neo UHPLC (Thermo Fisher Scientific, Waltham, MA USA). Trap and elute method using Thermo 300X5 Pepmap trap cartridge was used for trapping and IonOpticks Aurora 4 Ultimate XT 25×75 C18 column was used for separation. After trapping peptides were separated on a gradient from 92% A (water+0.1% FA)/8% B (80%ACN+20%Water+0.1% FA) to 45% B over 18 minutes. Spectrums were obtained using DIA method. Full scans from 380 to 980 m/z were acquired at a resolution of 240,000, 5ms max injection time and AGC target at 500%. For MS/MS isolation windows were set at 2m/z, injection time of 3ms and AGC target was set to 100% and collision energy was set 25%. Data were acquired using Xcalibur software (Thermo Scientific).

### MS data analysis

Data analysis of cell pellets was performed using the Fragpipe computational platform (V24.0) running the LFQ-MBR workflow with all default parameters^91–93^. Mass spectra were searched against a custom database containing the Chlamydomonas proteome (UP000006906) and *B. sola* MF6 target–decoy database. This database was supplemented with contaminants and decoys (50%). Mass spectra of supernatants were analyzed using DIA-NN 2.7.0^94^. Spectral library was searched against a custom database containing the Chlamydomonas proteome (UP000006906) and B. sola MF6 target–decoy database and data was analyzed with MBR option checked, MS1 accuracy set to 4 and MS2 at 10, quantification strategy QuantUMS^95^.

### Proteomic data analysis

Protein-level MaxLFQ intensities from the FragPipe v24.0 output were filtered to retain only bacterial protein groups (AC1V0C_#####) and log2 transformed (Table S8). Two of the 48 runs (34F7_C-_D2_1 and 34F7_C-_D3_1) were excluded as technical failures: both exhibited elevated contaminant spectra (27.84% and 19.54%), low spectral depth (0.10 and 0.14 times the cohort median total spectral count) and poor Pearson correlation to their respective replicates (r = 0.54 and 0.70).

### Differential expression of cell pellets proteomics data

Intensities were median-centered before testing, with the reference protein set differing between the two models. For the wild-type co-culture contrast, offsets were computed across all proteins quantified in the eight *Bso* MF6 day-2 runs. For the mutant including contrasts, offsets were computed on a core set, defined as protein groups quantified in all 23 day-2 runs and not encoded on chromosome 3, yielding 1,690 core proteins (Table S9). In both cases a per-run offset was calculated as the median log2 intensity across the reference set and applied to all quantified proteins in that run: intensities were centered by subtracting the offset and returned to the original scale by adding back the median of all offsets.

Differential abundance was assessed with limma 3.66.0^96^ in R 4.5.3, using lmFit on a means-parameterised design (∼ 0 + group) followed by contrasts.fit and eBayes with trend = TRUE and robust = TRUE^97^. A protein was considered quantifiable in a group when a MaxLFQ value was present in at least two of its replicates. Proteins encoded on chromosome 3 were excluded from all contrasts involving 27D6 or 34F7. Two models were fitted: the wild-type co-culture response (MF6 co-culture day 2 − MF6 monoculture day 2) was fitted on the eight *Bso* MF6 day-2 runs alone, and the six mutant contrasts (Table S10) were fitted in a single six-group model across all 23 day-2 runs. P values were adjusted within each contrast across all tested proteins using the Benjamini– Hochberg method, and proteins with an adjusted P < 0.05 were considered differentially abundant.

For the wild-type co-culture contrast only, proteins detected in one condition and not the other were additionally classified as ON/OFF. A protein was called ON in one condition and OFF in the other when it was present in at least three of the four replicates of the ON condition, absent from every replicate of the OFF condition, and its mean log2 intensity in the ON condition exceeded the median intensity of all proteins quantified in the OFF condition.

### Principal component analysis of cell pellets proteomics data

Proteins quantified in every run of a given set and encoded on chromosome 1 or 2 were retained, yielding 1,598 proteins across the 46 runs and 1,629 across the 24 co-culture runs. Their MaxLFQ intensities were log2 transformed and median-centred as described above. Each protein was then centred on its mean across runs, and principal components were obtained by singular value decomposition of the resulting run × protein matrix. The proportion of variance explained by each component was calculated from the squared singular values.

### Functional enrichment of the MF6 co-culture response

Enrichment analysis for the *Bso* MF6 co-culture contrast (MF6_C+_D2 versus MF6_C−_D2) was performed for COG functional categories and KEGG pathways using a one-sided Fisher’s exact test. The background comprised all proteins detected in this contrast that carried an annotation: 3,549 with an assigned COG category and 1,518 with an assigned KEGG pathway. Proteins annotated with more than one category contributed to each of them. Up- and down-regulated proteins were tested separately. The foreground comprised proteins passing an adjusted P < 0.05 with |log2 fold change| ≥ 1 in the differential abundance analysis, together with proteins receiving an ON call whose mean log2 intensity in the ON group exceeded the 50th percentile of the intensity distribution of the opposing group. KEGG pathways assigned to the Organismal Systems, Human Diseases and Drug Development classes, and to the Global and overview maps subclass, were excluded prior to testing. Terms were tested only when represented by at least five proteins in the background and two in the foreground, and P values were corrected by the Benjamini–Hochberg procedure within each ontology and direction (Table S11, day-2 co-culture DE).

### Clustering and functional enrichment of the core proteome

The 1,690 median-centred core proteins were z-score standardised across the 23 runs, and the dendrogram from Ward linkage on Euclidean distances was cut into eight clusters, a number chosen by inspection of the dendrogram to give interpretable groups without over-splitting. Runs were not clustered. Each cluster was tested for enrichment of COG functional categories and of KEGG pathways using a one-sided Fisher’s exact test, with the background restricted to core proteins carrying an annotation in the respective ontology. Proteins annotated with more than one category contributed to each of them. KEGG pathways assigned to the Organismal Systems, Human Diseases and Drug Development classes, and to the Global and overview maps subclass, were excluded prior to testing. Terms were tested only when represented by at least five proteins in the background and two in the cluster under test. P values were corrected by the Benjamini– Hochberg procedure within each cluster (Table S11, core expression cluster).

### Selection of candidate toxins and effectors

Every protein detected in the proteomics experiment was assessed against three independent criteria: extracellular localisation predicted by DeepLocPro, assignment of a Pfam accession labelled as toxin-related in the Toxinome classification^26^, and prediction as a T6SS effector (see below). Proteins meeting at least one criterion were retained, giving 369 candidates. Candidates were then filtered to those induced by the *Bso* MF6 in the presence of Chlamydomonas, defined from the day-2 *Bso* MF6 contrast (MF6_C+_D2 versus MF6_C−_D2) as either an adjusted P < 0.05 with positive log2 fold change, with no effect-size threshold applied, or an ON call in co-culture, leaving 34 proteins (Supplementary Table S14). This set was reviewed manually against the literature, domain architecture and genomic context to produce the short list discussed in the main text.

### Machine learning-based prediction of T6SS effector/toxin candidates

To identify candidate T6SS effector/toxin proteins, an XGBoost (eXtreme Gradient Boosting) classifier was trained. XGBoost builds an ensemble of decision trees sequentially, with each successive tree trained to correct the residual errors of its predecessors, resulting in a robust and accurate classifier. Hyperparameters were tuned to optimize performance and minimize false positives; the final model comprised 100 trees (n_estimators = 100), a maximum tree depth of 5 (max_depth = 5), and a minimum child weight of 7 (min_child_weight = 7). Both biochemical features (including mean hydrophobicity and sequence length) and genomic features (including distance from core T6SS genes and from known immunity genes) were integrated into the model, with all samples and corresponding features derived from the Integrated Microbial Genomes and Microbiomes (IMG) database^98^.

From the model output, only predictions with a model score > 0.75 that fell within eight genes upstream or downstream of core T6SS genes were retained. Candidates passing this filter were manually examined and prioritized for further validation; where two predictions were located in close proximity to the same VgrG protein, the one containing a known toxin domain was selected. For each candidate, model accuracy was further assessed by identifying additional protein domains using InterPro^79^ and evaluating structural similarity to known proteins using Foldseek^99^. To predict additional toxins across other bacterial species, Toxinome^26^, a lab-curated database compiling domains known to be toxic in bacteria was applied, and the model was run on Proteobacteria genomes predicted to encode a Type VI secretion system.

### Principal component analysis of the supernatant proteomics data

Bacterial proteins encoded on chromosomes 1 and 2 and quantified in at least three of four replicates in at least one group were retained, yielding 325 proteins. Missing values were imputed separately for each run from a down-shifted normal distribution of that run’s observed values (−1.8 SD mean; 0.3 SD width). Each protein was mean-centred across the 24 runs, and principal components were computed by singular value decomposition. Imputation was repeated 100 times, the sign of each component was aligned to the first draw, and the plotted scores and explained variance are the means over the 100 draws.

### Differential expression of supernatant proteomics data

Differential expression analysis of bacterial proteins was performed similar to whole cell proteomics. In short, proteins encoded on chromosome 3 were excluded from 27D6 or 34F7 contrasts and a protein was considered quantifiable in a group when it was present in at least two out of four replicates. A protein was called ON in one condition and OFF in the other when it was present in at least three of the four replicates of the ON condition, absent from every replicate of the OFF condition, and its mean log2 intensity in the ON condition exceeded the median intensity of all proteins quantified in the OFF condition.

A protein was considered MF6-specific if it was upregulated (log2FC ≥ 1, BH-adjusted p < 0.05) or ON (detected in ≥ 3 of 4 replicates with Chlamydomonas and 0 of 4 without) in the MF6 supernatant, and was neither upregulated nor ON in the 27D6 or 34F7 supernatants.

### Chlamydomonas RNA-seq

#### Algal growth, co-culture, and sample collection

*Chlamydomonas reinhardtii* CC-4403 (mt− gamete) was grown in 100 ml TAP medium in an Erlenmeyer flask on a shaker under continuous light for 2 days, with a 1:1 dilution after the first day. Cultures were then diluted to OD₇₅₀ = 0.5 in 10 ml TAP in a 50 ml Falcon tube. *Bso* MF6 was added to a final concentration of OD = 0.05, and a time 0 sample was harvested immediately. Additional samples were harvested every hour thereafter by centrifugation (2,000 rpm, 5 min), followed by flash-freezing in liquid nitrogen and storage at −80°C.

### RNA extraction, library preparation, and sequencing

Cell pellets were resuspended in 1 ml TRIzol and vortexed for 8 min at 4 °C. 200 µl chloroform was added, and samples were shaken for 60 s and centrifuged at 11,000 × g for 7 min. The upper aqueous phase was collected, diluted 1:1 with 100% ethanol, and RNA was purified using Direct-zol RNA MiniPrep Plus (Zymo research). RNA concentration was measured by NanoDrop (Thermo Scientific), and RNA integrity was assessed on an Agilent 2100 Bioanalyzer (RNA integrity number between 6 and 8).

Approximately 2 µg total RNA was used to prepare cDNA libraries using the TruSeq RNA kit (Illumina) according to the manufacturer’s instructions. Mean adapter-ligated cDNA size was 260 bp, as determined on a 2200 TapeStation (Agilent Technologies). Twelve barcoded cDNA libraries were pooled in equal proportions (by NanoDrop estimate) and sequenced in a single lane on a HiSeq2000 instrument (Illumina) by the Beijing Genomics Institute (Hong Kong), using 50-bp single-end reads.

### RNA differential expression analysis

Raw single-end reads were processed with fastp v1.0.1^100^ to remove adapters, low-quality bases (Q < 20; ≤20 % low-quality bases per read), poly-X tails, and reads shorter than 30 nt. Residual ribosomal RNA was removed using SortMeRNA v4.3.7^101^ against the Default database. Cleaned reads were aligned to the *Chlamydomonas reinhardtii* CC-4532 v6.1 genome^102^ with STAR v2.7.11b aligner^103^ with maximum intron length of 10,000. Gene-level counts were obtained with RSEM v1.3.3^104^ in reverse-strand mode. Genes with at least 10 reads in six or more libraries were retained for differential-expression analysis with DESeq2 v 1.46.0^105^. Each timepoint was contrasted against timepoint zero, with log₂ fold-change shrinkage applied. To capture time-course expression dynamics, the filtered count matrix was further modelled with ImpulseDE2 v1.11.0^106^ in single-condition mode, which classifies dynamically expressed genes as up- or down-regulated. Genes within each direction were soft-clustered into six groups by fuzzy *c*-means with Mfuzz v2.66^107^. GO Biological Process, Molecular Function and Cellular Component terms were tested for over-representation in each cluster with topGO v2.58.0^108^ using the *weight01* algorithm and Fisher’s exact test (node size ≥ 5, membership score ≥ 0.4), against the universe of all DESeq2-filtered genes. Terms supported by ≥3 genes at *P* < 0.01 were retained for visualisation. Principal-component analysis was performed on the top 1,000 most variable genes after *rlog* transformation.

### UV mutagenesis and screening of mutants for resistance phenotypes

To generate a UV mutant library, Chlamydomonas CC-4403 (mt⁻ gamete) culture was diluted to OD = 0.5. 780µl of this culture was spread onto each of 10 TAP agar plates and left to dry for ∼1 h in the dark. Once dry, plates were exposed to UV light for 1 min using a standard UV lamp in a biosafety cabinet, then immediately transferred to the dark and left overnight at room temperature. The following day, plates were moved to the light for an additional day of recovery. Cells were then scraped from the plates using 1 ml liquid TAP medium and spread onto new plates containing low-melting-point agarose embedded with *Bso* MF6 culture at a final OD = 0.02. Plates were incubated at 21°C under continuous light (∼50 µE). After approximately 1 week, surviving mutant patches appeared and were transferred to new agar plates. This was followed by several rounds of streak-purification on TAP medium supplemented with ampicillin (100 µg/ml), until four mutants appeared free of visible bacterial contamination. In a separate experiment, an existing UV mutant library (∼10,000 mutants) was replica-plated onto agar containing *Bso* MF6 (OD = 0.01), yielding 23 surviving mutants. Surviving colonies from both experiments were confirmed as resistant using the standard halo assay (described above). Two resistant mutants, one from each experiment, were selected for further analysis.

### Tetrad dissection of resistant UV mutants

Genetic crosses were performed following Dutcher (1995)^61^. Briefly, fresh mutant and wild-type strains of opposite mating type were each resuspended in 0.5 ml M-N/5 medium for ∼5 h to induce gametogenesis, then mixed together for ∼30 min to allow mating, spread onto agar plates, and kept in the dark for 5 days. After 5 days, vegetative cells were removed by scraping the plate surface with a flame-sterilized razor blade, and zygospores were collected into 100 µl double-distilled water using a flame-sterilized razor blade and spread onto a fresh agar plate for dissection.

Using a dissection microscope (SporePlay+, Singer Instruments, UK), zygospores were separated from any remaining vegetative cells and, the following day, individual tetrads were dissected into their four meiotic progeny. Each progeny clone was assayed by halo assay as described above, with six replicates per clone. Halo size was used to score progeny as either mutant-like (small halo) or wild-type-like (large halo), and 2:2 segregation of the resistance phenotype among tetrad progeny was used to confirm single-locus inheritance of the causative mutation.

### Bulked-Segregant-Analysis (BSA) of resistant UV mutants

This analysis was adapted from Tulin and Cross^42^. To prepare samples for sequencing, at least 10 wild-type-like and mutant-like segregants derived from backcrosses to strains M10 or P10 were isolated, grown at 21 °C,and pooled in equal proportions into 30 mL TAP cultures. Cells were resuspended in 8 mL of TEN buffer (10 mM Tris, 10 mM EDTA, 150 mM NaCl, pH 8) supplemented with 0.5% SDS and 0.1 mg/mL proteinase K, sonicated, incubated at 50 °C for 2 hours, and sonicated again to disrupt cellular integrity. The lysate underwent two rounds of extraction with 16 mL phenol-chloroform followed by precipitation with 100% ethanol. The resulting precipitate was rinsed with 70% ethanol, resuspended in TE buffer (10 mM Tris, 1 mM EDTA, pH 8), adjusted to 1 M NaCl, and subjected to a second phenol-chloroform extraction and ethanol precipitation. The final pellet was washed in 70% ethanol, resuspended in 200 μL TE, treated with 10 μg RNase A for 1 hour at 37 °C, and purified using the Qiaex gel extraction kit (Qiagen). Purified DNA was next applied to an Illumina PCR-free library preparation kit and sequenced using an Illumina NextSeq2000 machine. Resulting raw reads were quality-checked with FastQC v0.12.1^109^, and adapter sequences and low-quality bases were removed using fastp v1.0.1^100^. The trimming step included removal of poly-G tails and bases with a Phred quality score below 20 in more than 40% of the read, followed by discarding reads shorter than 35 bp. Trimmed reads were aligned to the *Chlamydomonas reinhardtii* CC-4532 v6.1 reference genome^110^ using BWA-MEM v0.7.19^86^ with default parameters. Alignments were converted to BAM, sorted and indexed with samtools v1.22^88^. SNPs and small indels were jointly called for the wild-type and mutant samples using FreeBayes v1.3.10^111^ under the *pooled-continuous* population model. To minimize low-quality variant calls, a minimum base quality of 20 and a minimum mapping quality of 30 were applied. The resulting VCF file was normalized using the bcftools v1.22^88^ *norm* command and variants located within repetitive or low-complexity genomic regions were removed. The remaining variants were annotated with SnpEff v5.2^112^ using a custom database built from the reference GFF annotation file. SNP-index values were calculated using the allele depth (AD) field for each variant, including only sites with a total read depth of at least 10. The SNP-index represents the fraction of reads supporting the alternate allele relative to the total reads at that position for a given sample. To identify variants associated with the phenotype, the SNP-index values of mutant samples were compared to those of the wild-type. Variants showing an absolute difference in SNP-index (ΔSNP-index) of 0.97 or greater were considered candidate mutations, indicating nearly complete fixation of the alternate allele in the mutant pool and its absence in the wild-type.

### *ZIP3* gene complementation

The complementation construct was generated by overlap PCR. The *ZIP3* gene, including 1 kb of upstream sequence, was amplified from genomic DNA using primers [Hyg_DNA1000 2.FOR GTATTTGAAGCGCTTTGCTGACCCTGGCCG; Hyg_DNA10002.REV GTGAGCATGCTTTTGAGGGCT], and the hygromycin B resistance cassette was amplified from plasmid pHyg3 using primers [Phyg_1000.FOR CTTTCTTGCGCTATGACACTTCC; Phyg_1000t 1.REV GGGTCAGCAAAGCGCTTCAAATACGCCCAGC]. Both products were gel-purified and used as templates for a second PCR joining the two fragments into a single construct, which was gel-purified and transformed into the mutants.

Transformation was performed as previously described^113^, with modifications. Cells were cultured to a density of approximately 8×10⁶ cells/mL, harvested, and resuspended in CHES buffer (10 mM N-cyclohexyl-2-aminoethanesulfonic acid, pH 9.25, 40 mM sucrose, 10 mM sorbitol) at room temperature to a final concentration of 8×10⁸ cells/mL. Electroporation was performed on 125 µL aliquots in 2 mm-gap cuvettes using a NEPA21 square-pulse electroporator (Nepa Gene, Japan). The pulse protocol consisted of two 8 ms poring pulses at 250 V and 150 V, followed by five 50 ms transfer pulses starting at 20 V with a 40% decay rate (sequentially decreasing to 20, 12, 7.2, 4.3, and 2.6 V). Immediately following electroporation, cells were transferred to 8 mL TAP medium supplemented with 40 mM sucrose. After overnight recovery at 24°C under low light, cells were collected by centrifugation and plated on TAP agar (1.5% w/v) containing 10 µg/mL hygromycin B.

### *zip3* and *cep12* liquid co-culture experiment

Relevant strains were grown overnight in liquid TAP in quadruplicate, then diluted to a final concentration OD=0.35. *Bso* MF6 was then added to a final concentration of OD=0.035. At 0, 24, and 48 h, a 1 ml sample was taken, centrifuged, and the pellet was snap-frozen and stored at −80°C. Chlorophyll was extracted using 0.5 ml 100% methanol. Following 5 min vortex and 12,000 g centrifugation, 200 µl was transferred to 96 well plate and absorbance was measured in 652 and 665 nm. Total chlorophyll concentration was defined by the equation 22.12*652+2.71*665.

### Annexin V/SYTOX Green PCD assay

Chlamydomonas was grown for 2 days in liquid TAP under continuous light (with one 1:1 back-dilution after 24 h). *Bso* MF6 was grown overnight at 28°C in liquid LB. For the experiment, triplicate 50 ml tubes containing 10 ml Chlamydomonas culture (final OD = 0.5 at 750 nm) were prepared, with or without *Bso* MF6 (final OD = 0.05 at 600 nm).

Samples were placed on a shaker at ∼150 rpm under continuous light. At the following timepoints post-inoculation, 1 ml of culture was transferred to an Eppendorf tube for staining: 0, 1, 2, 3, 4, and 5 h. Samples were centrifuged for 6 min at 3,000 x g and 15°C. The supernatant was discarded and samples were resuspended in 1 ml cold PBS; 0.5 ml was removed as an unstained control, and the remainder was centrifuged again and resuspended in 150 μl room-temperature ABB (10 mM HEPES, 140 mM NaCl, 2.5 mM CaCl2, pH 7.4) containing 500 nM SYTOX Green and Annexin V–Pacific Blue conjugate (1:20). Samples were incubated for 15 min at room temperature in the dark, after which 400 μl ABB was added to each sample before flow cytometry.

Excitation/emission wavelengths were as follows: Annexin V–Pacific Blue, 405/455 nm; chlorophyll, 635/730 nm; SYTOX Green, 488/523 nm. Flow cytometry was performed on a Cytek Aurora 4-laser spectral flow cytometer (Cytek Biosciences).

## Data analysis and availability

Proteomics data including the raw mass spectrometry files were submitted to ProteomeXchange Consortium *via* the PRIDE^114^ partner repository and can be accessed with the dataset identifiers PXD083366. *Bso* MF6 Tn5 mutant genome sequences, Chlamydomonas RNA seq data and *zip3-1*, *zip3-2* pooled mutant genome sequences are available in NCBI BioProject identifier PRJNA1517802. *Bso* MF6 new assembled genome is available in NCBI BioProject identifier GCA _056856995.2. The complete analysis pipelines are available at https://github.com/Michal-Breker-Lab/antagonistic-mechanisms-of-soil-microbiota; https://github.com/Michal-Breker-Lab/BSA-pipeline.

## Declaration of AI use in the manuscript preparation process

We used Claude.ai for data analysis and textual editing. Output was reviewed and edited by us.

## Acknowledgements

We thank Prof. Assaf Vardi (Weizmann Institute of Science) for generously providing the *cep12* mutant, for advice on PCD assays, and for helpful discussions and brainstorming. We thank Dr. Yael Friedmann (Bioimaging Unit, Silberman Institute of Life Sciences, Hebrew University of Jerusalem) for TEM imaging. M.B., E.M., and A.M. were funded by the European Research Council Starting Grant (ERC-StG), project Chloro-Import (grant no. 101117931). O.M.F was funded by the European Research Council Starting Grant (ERC-StG), project DryCoAdapt (grant no. 101077278) and Israel Science Foundation (ISF) (grant no. 2039/21). M.K. and O.M.F. were jointly funded by NSF-BSF grant (NSF award IOS #2421771 and BSF award #2024617). S.P. acknowledges funding from the Austrian Science Fund (FWF) (https://doi-org.uaccess.univie.ac.at/10.55776/COE7).

## Author contributions

Conceptualization, M.B. and O.M.F.; Methodology, M.B. and E.M. (algal screening platform and halo assays), O.M.F. and T.V. (ecological screens and halo assays), M.K. (proteomics), and A.L. (computational genome analysis); Software, A.M., M.A., and A.A.-A.; Formal analysis, A.M. (transposon-insertion mapping, proteome and transcriptome analysis, bulked segregant analysis, bacterial genomic analysis), M.A. (genome assembly, taxonomy and phylogenomic analyses), T.V. (ecological screens), A.A.-A. and A.L. (machine-learning analysis of bacterial genomes); Investigation, T.V. (isolate screen, halo assays, proteomics, TEM), E.M. (halo assays, algal genetics, transcriptomics, regulated cell death assays, TEM), T.S. (bacterial transposon library generation and screen) with valuable input from S.P., S.S. and M.K. (proteomics), O.K. (flow cytometry), and N.A.C. (halo assays); Resources, O.M.F.; Data curation, A.M.; Visualization, M.B., O.M.F., A.M., T.V., and E.M.; Writing – Original Draft, M.B., O.M.F., T.V., E.M., and A.M.; Writing – Review & Editing, all authors; Supervision, M.B., O.M.F., M.K., and A.L.; Project administration, M.B. and O.M.F.; Funding acquisition, M.B. and O.M.F.

---

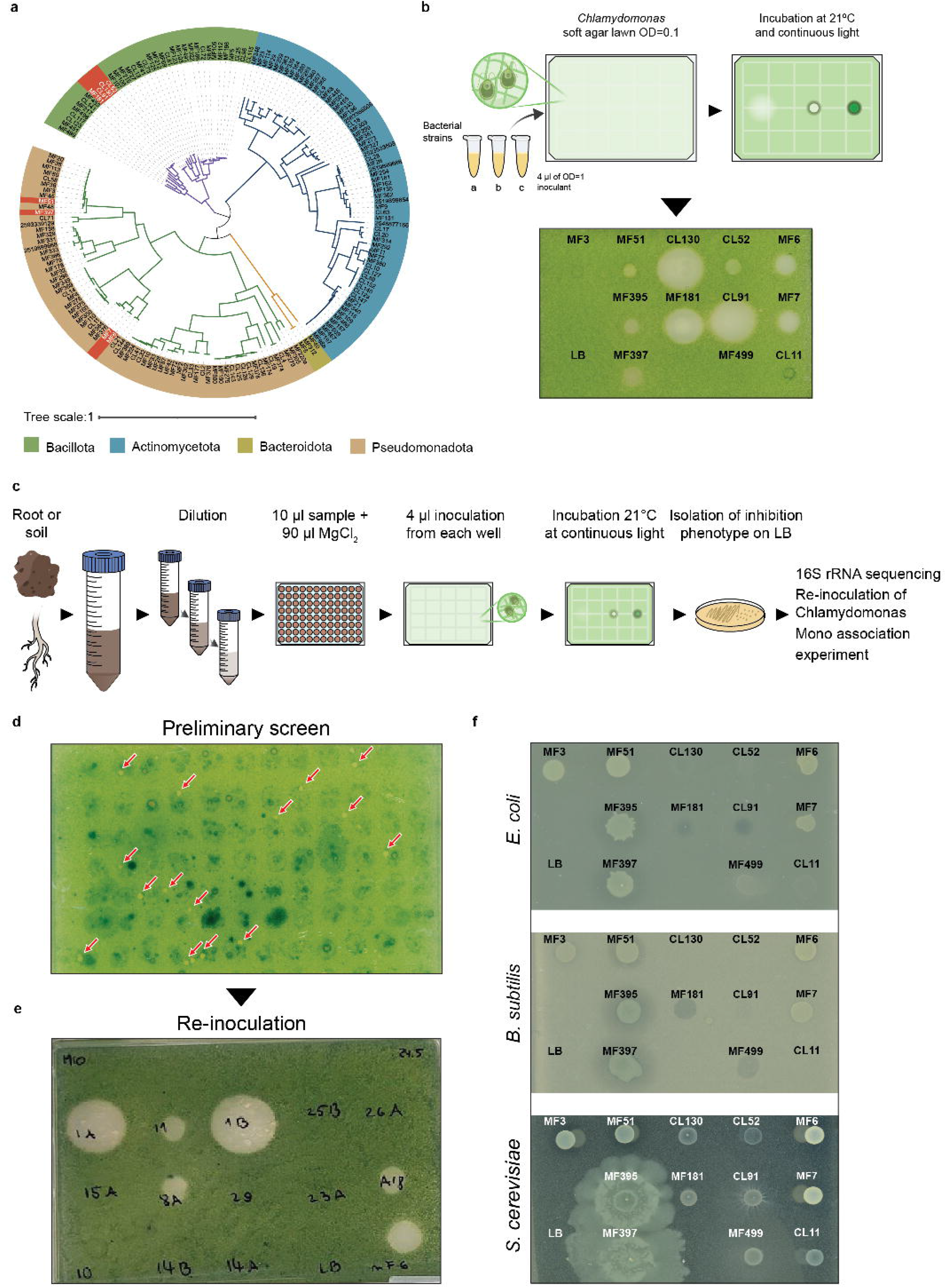

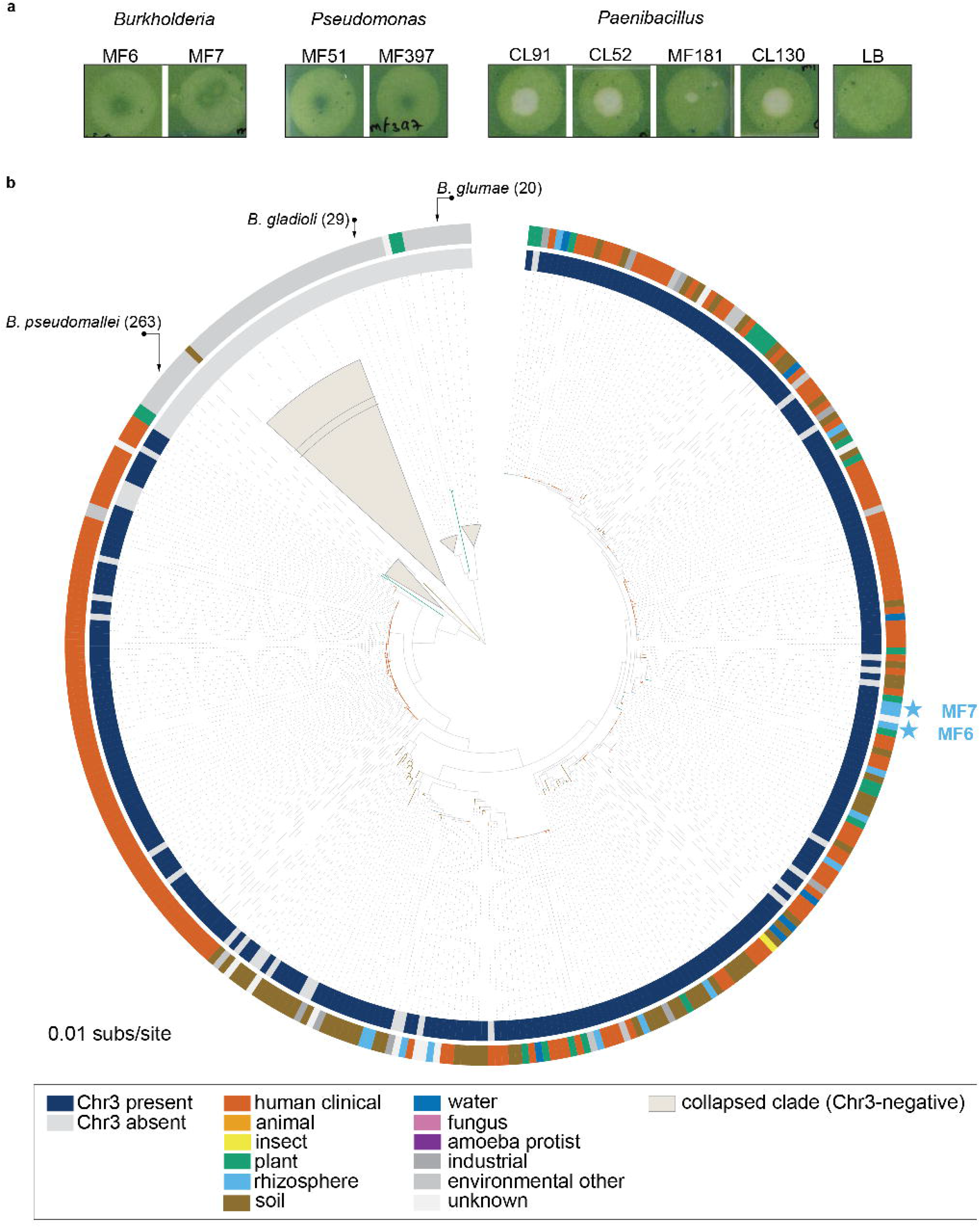

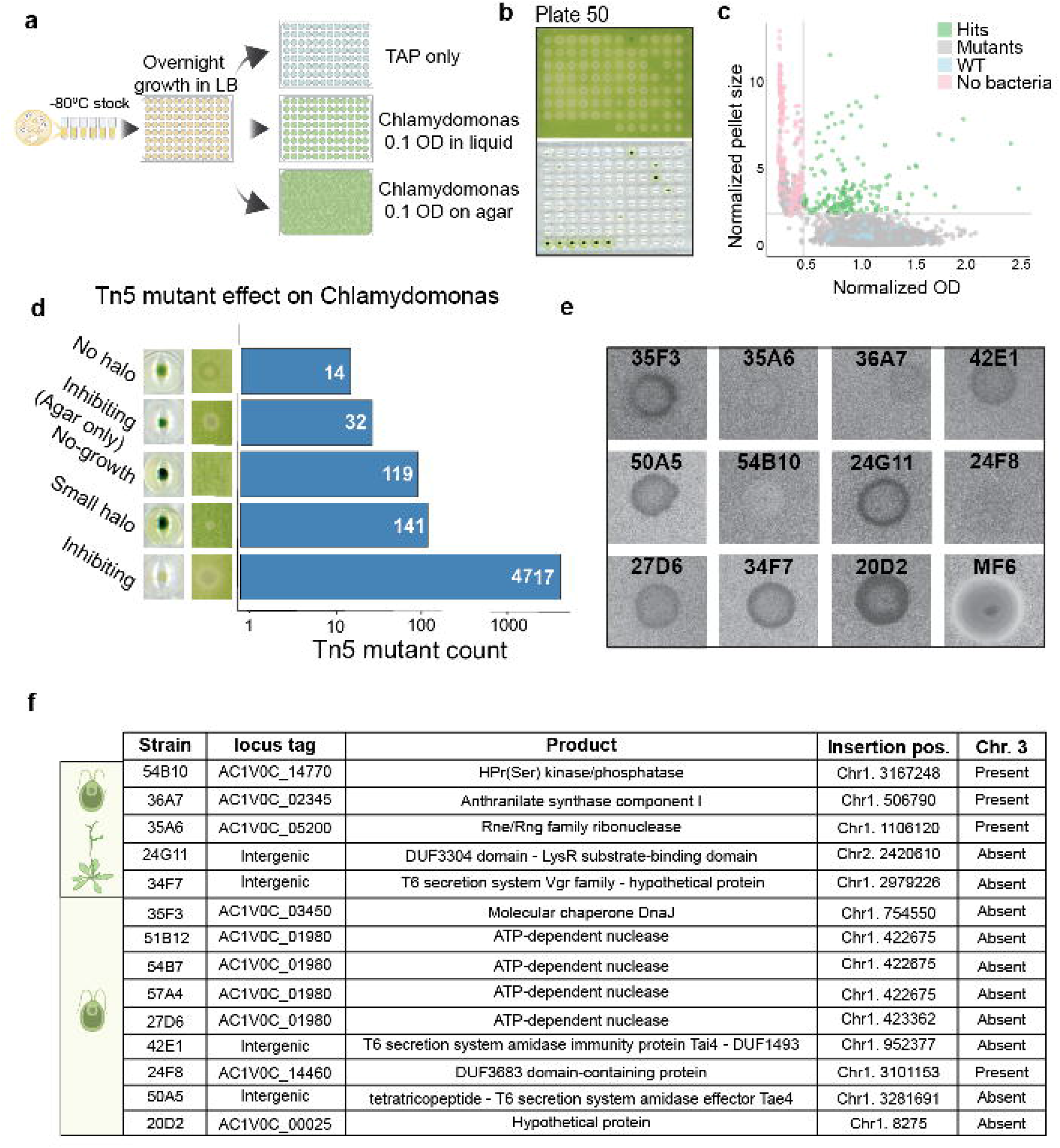

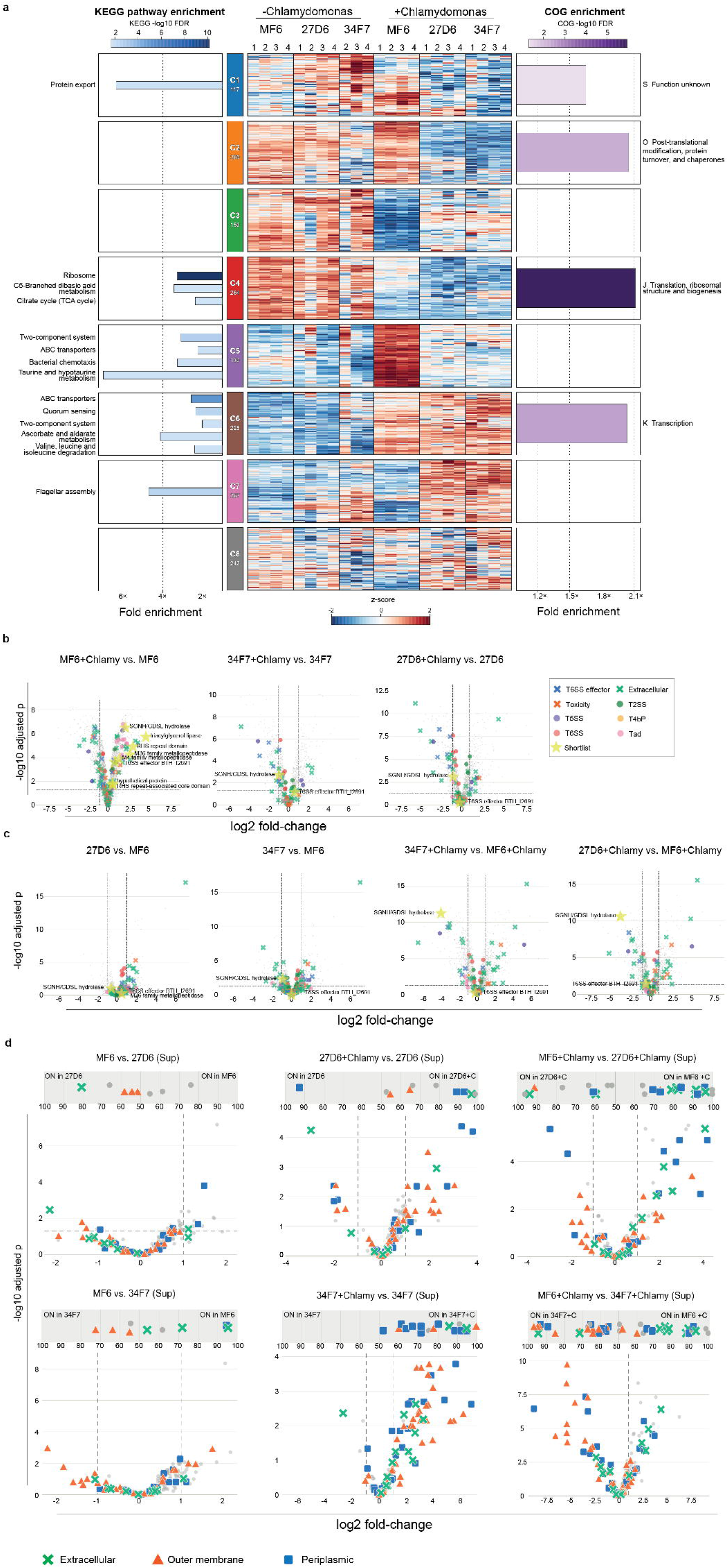

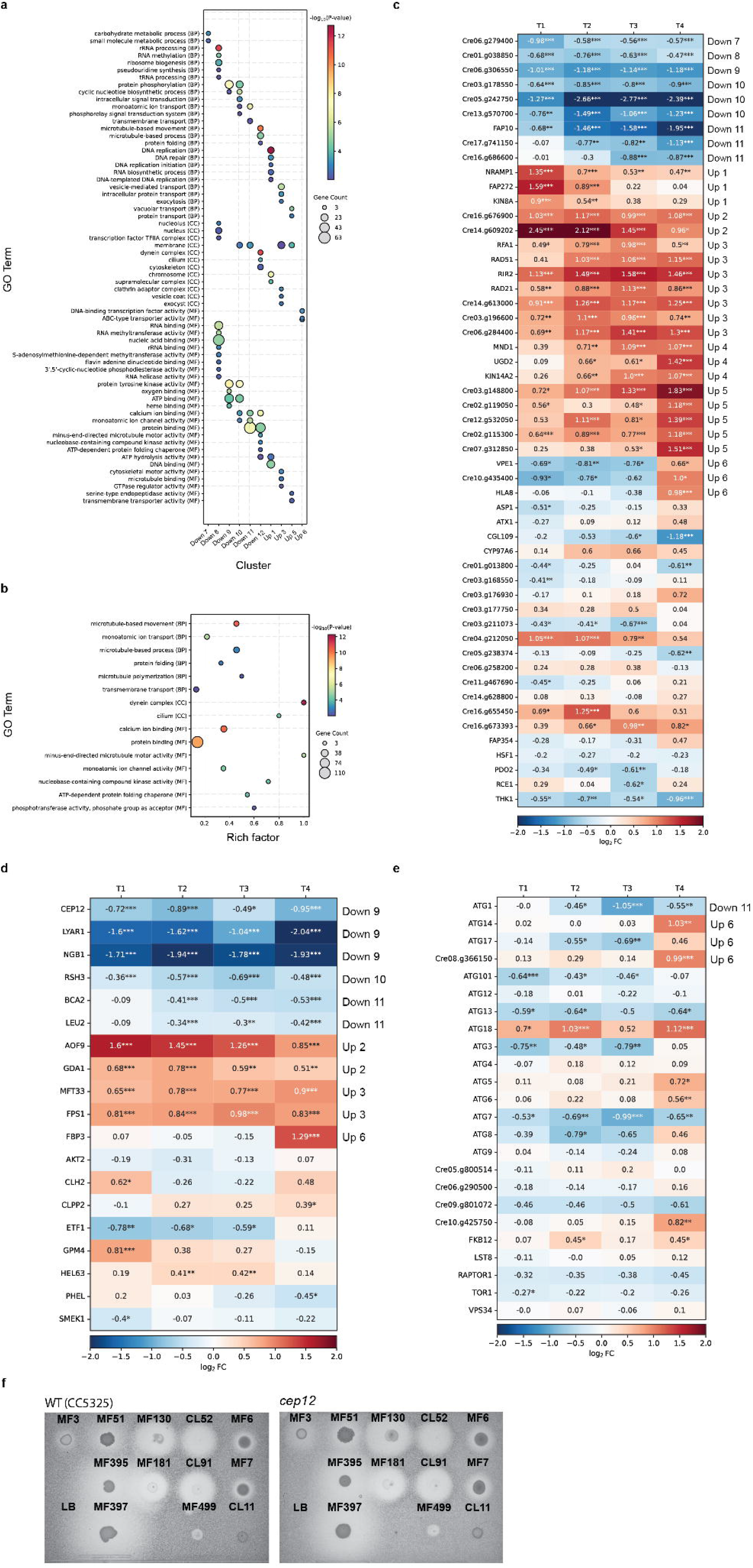

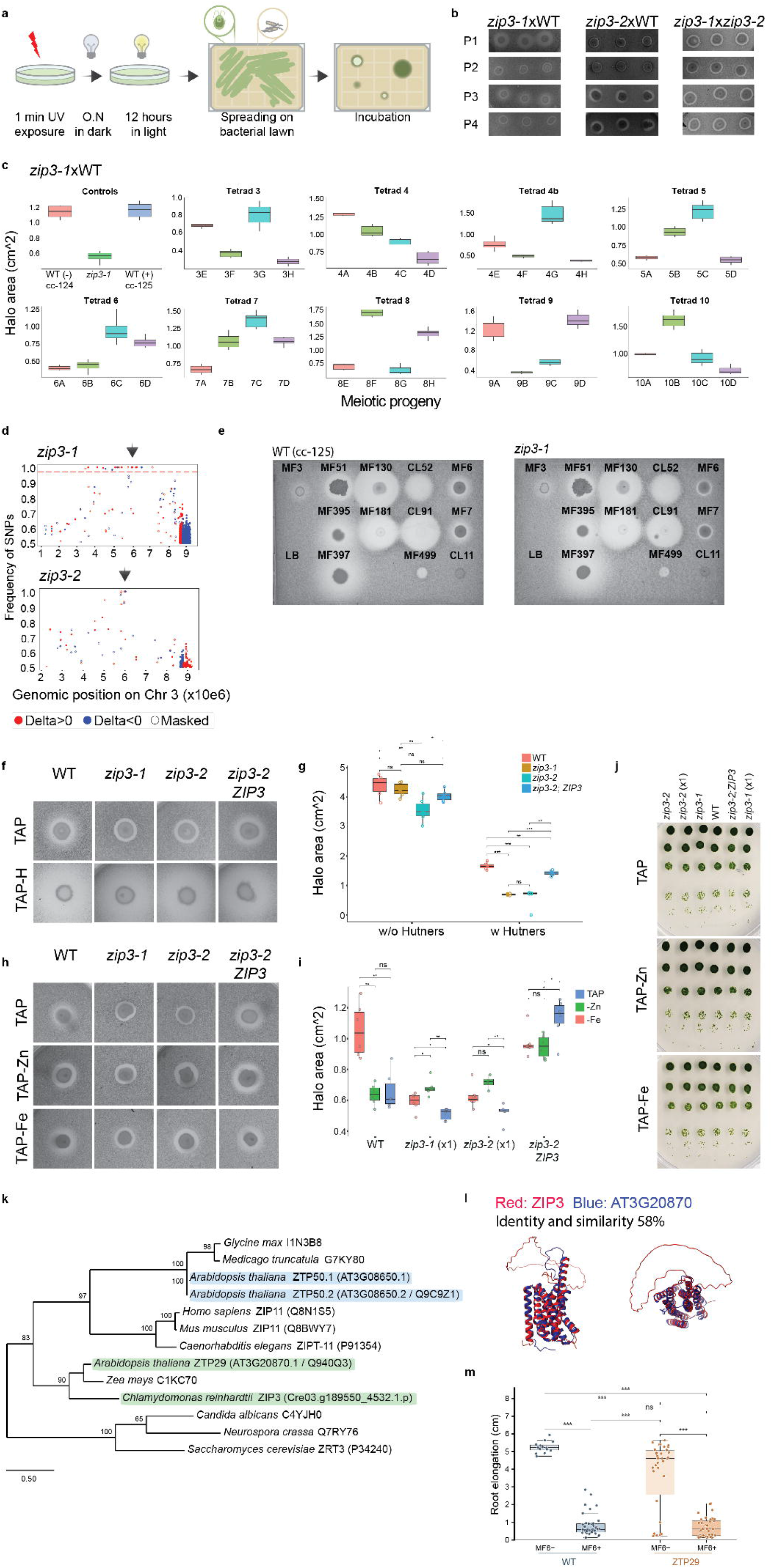

