## Supplemental information for "Conserved inhibitory mechanisms of root-associated bacteria towards *Arabidopsis thaliana* and *Chlamydomonas reinhardtii*"

**Legends for Extended Data Figures 1–6 and Supplementary Tables S1–S26**

**Extended Data Figure 1. Isolation and characterization of Chlamydomonas-inhibiting bacteria from a defined collection and from soil. Related to Fig. 1.** (a) Phylogenetic tree of the 149 bacterial strains isolated from *Arabidopsis thaliana* roots[1]. Tips are colored by phylum; strains that inhibited Chlamydomonas growth are highlighted in red. Adapted from Finkel et al, 2020[1]. (b) Schematic of the halo assay. Chlamydomonas was embedded in TAP soft agar (0.5% agar) at OD = 0.1, and 4 µl of each bacterial culture (OD = 1) was spotted onto the lawn. Plates were incubated at 21 °C under continuous light (50 µE) and scanned after 3–5 days. Below, a representative plate showing colorless zones of inhibition surrounding antagonistic strains; LB alone served as the negative control. (c) Schematic of the enrichment strategy for isolating Chlamydomonas-inhibiting bacteria directly from the environment. Soil or root suspensions were serially diluted in 10 mM $MgCl_2$, spotted onto Chlamydomonas soft-agar (0.5% agar) lawns, and incubated at 21 °C under continuous light. Bacteria were recovered from within the resulting halos, purified on LB, identified by 16S rRNA gene sequencing, and re-tested on Chlamydomonas and in Arabidopsis mono-association experiments. (d) Representative preliminary screen of a soil dilution series spotted onto a Chlamydomonas lawn. Arrows indicate colonies surrounded by zones of inhibition, which were picked for purification. (e) Re-inoculation of the purified isolates onto fresh Chlamydomonas lawns to confirm halo formation in pure culture. (f) Target range of the Chlamydomonas-inhibiting strains against *Escherichia coli* (top), *Bacillus subtilis* (middle), and *Saccharomyces cerevisiae* (bottom). Strains were spotted onto lawns of each target organism embedded in soft agar and imaged after 1–2 days at 28 °C. LB alone served as the negative control.

**Extended Data Figure 2. The requirement for contact differs among inhibitory genera, and chromosome 3 is unevenly distributed across *Burkholderia*. Related to Fig. 2.** (a) Halo formation by taxonomically diverse bacterial isolates on Chlamydomonas lawns. Autoclaved cellulose filters (25 mm diameter, 0.22 μm pore size) were placed onto soft-agar lawns of Chlamydomonas, separating the bacteria from the algal cells while permitting diffusion of soluble molecules. Overnight cultures of each isolate were normalized to $OD_{600}$ = 1 and 4 μl was spotted onto each filter; 4 μl of sterile LB was spotted as a medium-only control (rightmost panel). Plates were incubated for 5 days under standard Chlamydomonas growth conditions. Isolates are grouped by genus: *Burkholderia* (MF6, MF7), *Pseudomonas* (MF51, MF397) and *Paenibacillus* (CL91, CL52, MF181, CL130). Representative plates are shown. (b) Distribution of Chromosome 3 across the genus-level phylogeny of *Burkholderia* sensu lato. Maximum-likelihood phylogeny of 763 complete/chromosome-level *Burkholderia* sensu lato genomes with available isolation-source metadata, based on 120 GTDB single-copy marker genes (5,010 masked amino-acid columns; IQ-TREE, LG+F+R4, 1,000 ultrafast bootstrap replicates). Thirty-seven isogenic transposon mutants of *B. sola* R-12632 (PRJEB40633) were excluded, retaining only the parental strain; no other dereplication was performed. The tree is rooted on the ingroup stem, with a 123-genome outgroup and three near-clonal blocks (312 genomes) collapsed for display. Branches are colored by isolation source where all descendants agree; disagreeing

branches are marked separately. Chromosome 3 presence/absence is shown per tip (present in 282/763 genomes, 37%). *Bso* MF6 and MF7 are indicated. Collapsed chromosome 3-negative clades are labeled with genome counts (*Paraburkholderia*, *n* = 81; *Caballeronia*, *n* = 14; *Mycetohabitans*, *n* = 13; *Burkholderia*, *n* = 9; *B. glumae*, *n* = 20; *B. gladioli*, *n* = 29; *B. pseudomallei*, *n* = 263).
Only 444/763 genomes have a unique marker sequence; the remaining 393 (52%) are identical to at least one other genome (largest group, *n* = 70). As IQ-TREE cannot output true zero-length branches (floor: $2.55 \times 10^{-6}$ substitutions/site), these near-identical genomes appear fully resolved despite lacking real phylogenetic signal: splits not recovered by an independent FastTree analysis of the same dataset had a median bootstrap of 19, versus 95 for splits both methods recovered. Relationships among closely related strains are resolved using ANI analysis.

**Extended Data Figure 3. A Tn5 transposon screen identifies *Bso* MF6 loci required for inhibition of Chlamydomonas. Related to Fig. 3.** (a) Schematic of the transposon mutant library screen. Individual mutants were recovered from frozen glycerol stocks, grown overnight in LB, and replica-plated in parallel onto TAP-only control plates, into Chlamydomonas liquid co-culture (0.1 OD), and onto Chlamydomonas soft-agar (0.5% agar) lawns (0.1 OD). (b) Representative primary screening plate (Plate 50), showing the agar halo assay (Top) and liquid co-culture pellet assay (Bottom) used to score each arrayed mutant. (c) Normalized Chlamydomonas pellet size is plotted against normalized bacterial OD when grown alone in TAP; dashed lines indicate the thresholds used to classify hits (green) relative to the bulk mutant library (grey), wild-type *Bso* MF6 (blue), and Chlamydomonas-only controls with no added bacteria (pink). (d) Classification of the 5,023 screened Tn5 mutants into five phenotypic categories based on the combined agar and liquid assay readouts, with representative images for each: no halo (14), inhibiting on agar only (32), no bacterial growth (119), small halo (141), and inhibiting (4,717). (e) Representative halo assays of 11 candidate non-inhibitory mutants that passed tertiary validation, alongside wild-type *Bso* MF6. (f) Genomic identity of the 14 Tn5 insertions: locus tag, predicted protein product, insertion coordinate, and whether chromosome 3 is retained (Present) or lost (Absent) in that mutant. Bottom block, mutants that lost the ability to inhibit Chlamydomonas only; Top block, mutants that additionally lost the ability to inhibit Arabidopsis.

**Extended Data Figure 4. Proteome and secretome remodelling of *Bso* MF6 and its 27D6 and 34F7 mutants during co-culture with Chlamydomonas. Related to Fig. 3.** (a) KEGG pathway (left) and COG category (right) enrichment for each of the eight protein co-abundance clusters (C1–C8; number of proteins per cluster given below) identified in the core bacterial proteome (1690 proteins that were detected in all samples) at day 2 of co-culture with Chlamydomonas (FDR < 0.05). Center, standardized (z-score) protein abundance across genotype, condition (± Chlamydomonas), and biological replicate (1–4; replicate 1 of 34F7 monoculture excluded as a technical outlier, see Online Methods). (b) Volcano plots of differential protein abundance in co-culture with Chlamydomonas versus monoculture, for wild-type *Bso* MF6, 34F7, and 27D6. Dashed lines mark $|\log_2 FC| = 1$ and the significance threshold; selected secretion- and toxin-associated proteins are labeled. (c) Volcano plots of differential protein abundance between genotypes. 27D6 versus wild-type and 34F7 versus wild-type, in monoculture (left two panels) and in co-culture with Chlamydomonas (right two panels). (d) Volcano plots of differential protein abundance in the culture supernatant (Sup) at day 2, comparing wild-type *Bso* MF6 with the 27D6 (top row) and 34F7 (bottom row) mutants. Left, wild-type versus mutant in monoculture. Middle, mutant in co-culture with Chlamydomonas (+Chlamy) versus monoculture. Right, wild-type versus mutant in co-culture. Positive $\log_2$FC indicates higher abundance in the first-named condition. Dashed lines mark $|\log_2 FC| = 1$ and the significance threshold. Proteins are shaped and colored by subcellular localization predicted with DeepLocPro: extracellular (green crosses), outer membrane (orange triangles) and periplasmic (blue squares); all other proteins are shown in grey. Strips above each plot show proteins detected in only one of the two compared groups (ON/OFF), for which fold change cannot be calculated. Position along the strip indicates the percentile of the group the protein is missing from (50 at the centre, 100 at each edge), and the group in which the protein is detected ("ON in…") is labeled on the corresponding side.

**Extended Data Figure 5. GO enrichment and candidate-gene expression support acute deflagellation and a non-canonical, regulated death program during infection. Related to Fig. 5.** (a) Gene Ontology (GO) term enrichment (Biological Process, Cellular Component, and Molecular Function) across the significant temporal expression clusters identified during infection (see Fig. 5a, b). Dot size indicates gene count per term; color indicates $-\log_{10}$($p$-value) (topGO weight01 algorithm, Fisher's exact test). (b) GO term enrichment for the merged acute-response gene set (clusters 1 and 12), plotted by Rich factor (=Fraction of all genes annotated with a given term that fall within the cluster). Dot size and color as in (a). (c) Expression ($\log_2$ fold-change relative to T0) at T1–T4 post-infection of programmed cell death–associated genes, manually curated from three published studies of PCD[2–4]. Cluster assignment is indicated at right for genes belonging to a temporal cluster; the remaining genes did not meet cluster-membership criteria. Asterisks denote DESeq2 adjusted-$p$ significance (*$p < 0.05$, **$p < 0.01$, ***$p < 0.001$). (d) Expression of candidate programmed cell death (PCD)-related genes, selected based on homology to conserved algal cell-death genes identified in Mizrachi et al.[5], across infection. Cluster assignment (where applicable) is indicated at right; asterisks as in (c). (e) Expression of core autophagy (ATG) machinery and regulatory genes across infection. Cluster assignment (where applicable) is indicated at right; asterisks as in (c). (f) Halo assays of the parental strain CC-5325 (top) and the *cep12* insertional mutant (bottom) against a panel of twelve bacterial isolates; quantified in Fig. 5i.

**Extended Data Figure 6. Genetic mapping and functional validation identify ZIP3 as a zinc/iron transporter conferring metal-dependent susceptibility to *Burkholderia*. Related to Fig. 6.** (a) Schematic of the UV mutagenesis and selection scheme used to *Bso* MF6-resistant Chlamydomonas mutants: cells were UV-irradiated (1 min), recovered overnight in the dark, given 12 h in the light, then spread onto a *Bso* MF6 soft-agar (0.5% agar) lawn; resistant colonies were picked after incubation.(b)

Representative halo assays of the four meiotic progeny (P1–P4) of a single representative tetrad from each of the *zip3-1* × wild-type and *zip3-2* × wild-type backcrosses, each progeny spotted with *Bso* MF6 in triplicate, showing 2:2 segregation of the resistance phenotype. Right, the four progeny (P1–P4, triplicate spots) of a single representative tetrad from a *zip3-1* × *zip3-2* intercross; all four retained the resistant phenotype, consistent with non-complementation. (c) Halo area ($cm^2$) of individual meiotic progeny from nine dissected tetrads of a *zip3-1* × wild-type cross, alongside wild-type (cc-124, cc-125) and *zip3-1* controls, confirming 2:2 segregation of the resistance phenotype. Boxes show median and interquartile range. (d) Bulked-segregant analysis (BSA) of *zip3-1* and *zip3-2* resistant progeny pools. SNP frequency is plotted against genomic position on chromosome 3; arrows indicate the mapped peak, coincident with the ZIP3 locus in both mutants. Red, $\Delta$(SNP-index) $> 0$; blue, $\Delta$(SNP-index) $< 0$; open circles, masked calls; dashed line, the 0.97 $\Delta$(SNP-index) threshold used to call candidate variants (see Online Methods). (e) Halo assays of the parental strain cc-125 (top) and *zip3-1* (bottom) against a panel of twelve bacterial isolates; quantified in Fig. 6d. (f) Representative halo assays of wild-type, *zip3-1*, *zip3-2*, and the *zip3-2*;ZIP3 complemented line on TAP and TAP lacking Hutner's trace elements (TAP-H). (g) Quantification of halo area ($cm^2$) for the genotypes and conditions in (f) [unpaired t.test followed by BH correction $n$ = 4-6]. (h) Representative halo assays of wild-type, *zip3-1*, *zip3-2*, and *zip3-2*;ZIP3 on TAP, TAP lacking zinc (TAP-Zn), and TAP lacking iron (TAP-Fe). (i) Quantification of halo area ($cm^2$) for the genotypes and conditions in (h). [unpaired t.test followed by BH correction $n$ = 4-6]. (j) Serial-dilution growth assays of the indicated genotypes on TAP, TAP-Zn, and TAP-Fe in the absence of bacteria, confirming that zinc or iron omission alone does not impair growth. (k) Maximum-likelihood phylogenetic tree of ZIP-family zinc/iron transporters across eukaryotes, rooted with fungal ZRT3 orthologs. Node labels indicate bootstrap support; ZIP3 and its closest plant homolog, ZTP29, are highlighted in green, and the plasma-membrane-type ZTP50 paralogs in blue. Scale bar, substitutions per site. (l) Predicted structural alignment of Chlamydomonas ZIP3 (red) and Arabidopsis ZTP29/AT3G20870 (blue), shown from two angles, with RMSD and sequence identity/similarity indicated. (m) *ztp29* does not protect Arabidopsis from MF6-mediated suppression of root elongation. Root elongation of wild-type and *ztp29* seedlings grown without (MF6−) or with (MF6+) *Bso* MF6. Boxes show the median and interquartile range, whiskers extend to the most extreme value within 1.5 × IQR, and points show individual seedlings ($n$ = 15, 38, 34 and 36 from left to right). Data were analyzed by two-way ANOVA (genotype, $p = 0.0037$; MF6, $p = 2.5 \times 10^{-32}$; genotype × MF6, $p = 9.5 \times 10^{-4}$) followed by Tukey's HSD; ***$p < 0.001$, ns, not significant. MF6 suppressed root elongation in both genotypes, and infected wild-type and *ztp29* seedlings were indistinguishable, indicating that loss of ZTP29 confers no protection. See also Online Methods.

### Supplemental tables legends

**Supplementary Table S1. Bacterial isolates used in this study. Related to Fig. 1.** The 149 bacterial isolates used in this study, drawn from the 185-member synthetic community of genome-sequenced strains isolated from surface-sterilized Arabidopsis roots described in Finkel et al. (2020)[1]. *Strain* gives the isolate identifier used in that collection and throughout this manuscript (108 MF-series and 41 CL-series isolates); *Name* gives the organism name and strain designation as deposited. *NCBI Assembly or BioProject accession (GenBank)* and *NCBI Assembly Accession (RefSeq)* give the corresponding genome accessions where a public assembly exists; for one isolate (CL004) only a BioProject accession is available. *IMG Genome ID* gives the identifier of the sequenced genome in the JGI Integrated Microbial Genomes database, available for all 149 isolates. NA indicates that no corresponding accession exists. 26 isolates have IMG genomes with no corresponding NCBI assembly.

**Supplementary Table S2. Chlamydomonas halo assay results upon infection with the Arabidopsis root bacterial collection, with comparative Arabidopsis primary root elongation data from Finkel et al.[1]. Related to Fig. 1.** Halo assay: Taxonomy, halo formation (Y/N) on Chlamydomonas soft-agar lawns, and growth on TAP medium alone in the absence of an algal lawn (scored as 0 = no visible growth, 0.5 = weak or partial growth, 1 = robust growth) for each strain in the Arabidopsis root bacterial collection. Finkel et al. 2020: Per-seedling primary root elongation measurements (cm) for the overlapping bacterial collection on Arabidopsis, reproduced from Finkel et al[1], used for the comparison in Fig. 1b. Statistics were determined separately for each independent experiment (mono1 and mono2) using Welch's t-test with Benjamini–Hochberg correction for multiple comparisons ($q < 0.05$). Data from additional experiments are included but were not analyzed statistically, as they represent partial subsets of the collection. Inhibition towards Arabidopsis was defined as a significant reduction in at least one of two experiments together with a mean reduction in root elongation of at least 25%.

**Supplementary Table S3. Primary root elongation of Arabidopsis seedlings inoculated with pine-forest soil bacterial isolates. Related to Fig. 1.** Per-seedling primary root elongation (cm) 12 days after transfer onto plates inoculated with bacteria isolated from pine-forest soil using Chlamydomonas halo formation as the selection criterion (see Extended Data Fig. 1c–e), or $MgCl_2$ buffer alone (control). "Inhibiting Chlamydomonas" indicates whether the isolate formed a halo on Chlamydomonas lawns in a secondary screening.

**Supplementary Table S4. Leaf area of Arabidopsis seedlings inoculated with pine-forest soil bacterial isolates. Related to Fig. 1.** Per-seedling leaf area (cm²) 12 days after transfer onto plates inoculated with bacteria isolated from pine-forest soil using Chlamydomonas halo formation as the selection criterion (see Extended Data Fig. 1c–e), or $MgCl_2$ buffer alone (control), across two independent experimental replicates (rep 1, rep 2). "Inhibiting Chlamydomonas" indicates whether the isolate formed a halo on Chlamydomonas lawns; values match Supplementary Table S3.

**Supplementary Table S5. Taxonomic classification of pine-forest soil- and rhizosphere-derived bacterial isolates. Related to Fig. 1.** 16S rRNA gene-based taxonomic classification (phylum through genus) of bacterial isolates recovered from pine-forest soil and rhizosphere samples using Chlamydomonas halo formation as the enrichment criterion (see Extended Data Fig. 1c–e). "Chlamydomonas inhibited" indicates halo formation on Chlamydomonas lawns; isolate codes correspond to those used in Supplementary Tables S3 and S4.

**Supplementary Table S6. Secretion system, pilus, and flagellar gene content of *Bso* MF6 and seven related bacterial genomes. Related to Fig. 2.** TXSScan (MacSyFinder) hits for secretion-system, pilus, and flagellar gene models across the genomes of *Bso* MF6 and seven related strains, one row per detected protein per system instance. The model_fqn field reads "diderm" on every row because 17 of TXSScan's 19 available models fall under that heading. This reflects TXSScan's model organization, not a claim about the organisms' cell envelope type; no monoderm-specific model was searched. Type VII secretion (T7SS), shown separately in Fig. 2a, was called by manual criteria (see Online Methods). system_wholeness and system_score reflect MacSyFinder's model-completeness metrics for each system instance; hit_status indicates whether a gene is a mandatory, accessory, or neutral component of that model.

**Supplementary Table S7. Genome metadata for the *Burkholderia* phylogeny. Related to Extended Data Fig. 2.** Metadata for the 763 genomes included as tips in the chromosome-1 core-genome phylogeny, one row per genome. *Accession* gives the NCBI assembly accession; *Organism_name*, *Genus*, *Species* and *Strain* give the NCBI taxonomic assignment as submitted; *Assembly_level* gives the assembly status reported by NCBI. *Host_category* is a harmonized isolation-source category (human_clinical, soil, plant, rhizosphere, water, animal, insect, fungus, amoeba_protist, industrial, environmental_other, or unknown) assigned from the verbatim BioSample fields reproduced in *Raw_host*, *Raw_isolation_source* and *Raw_host_disease*; *Host_evidence* records which of these fields the assignment was based on, or notes that no usable metadata were available. *Geo_loc_name* gives the reported country and, where available, region of isolation. *Chr3_present* indicates presence or absence of chromosome 3. The set comprises 646 *Burkholderia*, 84 *Paraburkholderia*, 16 *Caballeronia*, 14 *Mycetohabitans* and 3 *Trinickia* genomes across 80 species-level labels (68 genomes are assigned only to "sp."), with *B. pseudomallei* (197), *B. multivorans* (54) and *B. glumae* (52) the most represented. By origin, 276 genomes are human clinical isolates, 161 from soil and 86 plant-associated, while 136 lack usable isolation metadata; chromosome 3 is present in 252 of the 763 genomes. The final row is strain *Bso* MF6, a laboratory isolate not deposited in NCBI, for which metadata were curated manually. Empty cells indicate fields absent from the source record.

**Supplementary Table S8. Raw $\log_2$ MaxLFQ intensities per protein per proteomic run for *Bso* MF6, 27D6, and 34F7. Related to Fig. 2**. $\text{Log}_2$-transformed raw MaxLFQ intensities for each detected bacterial protein across all 48 proteomic runs (genotype [27D6/34F7/MF6] × condition [C−, alone; C+, with Chlamydomonas] × day [D2, D3] × biological replicate [1–4]; column headers follow the format genotype_condition_day_replicate). See Table S9 for normalized values used in differential abundance analysis. Blank cells indicate the protein was not quantified in that run. Includes the two runs (34F7_C-_D2_1, 34F7_C-_D3_1) excluded from all downstream analyses as technical outliers (see Online Methods); protein-level values may still be present in these columns even though the run itself was excluded.

**Supplementary Table S9. Core-set-centered $\log_2$ MaxLFQ intensities per protein, day 2 proteomic runs. Related to Fig. 2**. $\text{Log}_2$ MaxLFQ intensities from Table S8, each run centered on its own median across a defined core set of proteins (in_core_set = "yes," 1,690 of 4,023 proteins), rendering values comparable across runs. This is the matrix used for differential abundance modeling (Fig. 2b, c). Day 2 only, therefore no equivalent centered matrix exists for Day 3. 23 runs (27D6, 34F7, MF6 × C−/C+ × 4 replicates), excluding the QC-failed run 34F7_C-_D2_1.

**Supplementary Table S10. Differential protein abundance and presence/absence calls across seven day-2 *Bso* MF6 proteomic contrasts. Related to Fig. 2 and Extended Data Fig. 4.** Long-format differential abundance results, one row per protein per contrast, for the seven pairwise comparisons tested at day 2 (*contrast* column; group_A vs. group_B). *logFC* is group_A minus group_B, with accompanying *p_value* and Benjamini–Hochberg-adjusted *p_adj* (limma moderated *t*-test); *n_detected_A* and *n_detected_B* give the number of replicates in which the protein was quantified in each group. *ON_group* indicates the group where protein is treated as ON and *ON_percentile* indicates its LFQ percentile in the opposite group. *DeepLocPro_localisation* gives predicted subcellular localization; *toxin_Pfam_domains* flags matches to toxin-associated Pfam domains; *T6SS_effector_probability* is the score from the XGBoost T6SS effector classifier (see Online Methods). The final five columns report biosynthetic gene cluster (BGC) membership predicted by antiSMASH: *antiSMASH region* gives the identifier of the BGC region containing the encoding gene (23 regions across the three replicons); *BGC class* gives the predicted product class of that region; *antiSMASH role* gives the gene's assigned role within the cluster (core biosynthetic, additional biosynthetic, transport-related, regulatory, or other), left blank for cluster genes without an assigned role; *MIBiG accession* and *MIBiG similarity* give the closest known cluster in the MIBiG database and the corresponding similarity score. Of the 3,935 proteins in the table, 293 fall within a predicted BGC region and 153 carry an assigned role; all five BGC fields are blank for proteins outside predicted regions.

**Supplementary Table S11. COG and KEGG enrichment for the day-2 *Bso* MF6 co-culture proteomic response and the eight core expression clusters. Related to Fig. 2 and Extended Data Fig. 4.** Fisher's exact test enrichment results (Benjamini-Hochberg-corrected) for two independent analyses, distinguished by the analysis column. "Day-2 co-culture DE" tests proteins significantly more or less abundant in *Bso* MF6 co-cultured with Chlamydomonas versus monoculture at day 2 (direction; set = the single contrast MF6_C+_D2_vs_MF6_C-_D2), against the detected bacterial proteome as background. "Core expression cluster" tests each of the eight co-abundance clusters (set = cluster 1–8) against the median-centered core proteome. Within each test, *k* is the number of query-set proteins carrying the term and *n* is the size of the query set; *K* and *N* are the corresponding counts for the background. Query set and background are both restricted to proteins annotated in the tested ontology (ontology column), so *n* and

*N* are smaller than the full set sizes. “fold_enrichment” calculated as (*k/n*)/(*K/N*). The final column lists the identifiers of the proteins carrying the term, pipe-delimited.

**Supplementary Table S12. Transposon insertion site details for sequenced *Bso* MF6 mutant clones. Related to Fig. 3.** Insertion-site mapping for 16 *Bso* MF6 transposon mutant clones (16 sequenced isolates total; clones 35A6 and 36A7 each represented by two independently isolated and sequenced colonies), determined by Illumina or Nanopore sequencing (Platform) as indicated per clone. tsd_len_junction and tsd_len_insert give the target-site duplication length measured from the junction reads and, where available (Nanopore only), from the reads spanning the whole cassette. Cassette_orientation indicates the orientation of the transposon relative to the reference genome. For insertions falling within a coding sequence, Locus_tag, Gene, and Product identify the disrupted gene, with Cds_start/Cds_end/Cds_strand/Cds_len_bp giving its genomic coordinates and bp_into_cds/Codon/aa_pos_of locating the insertion within the coding sequence and predicted protein.

**Supplementary Table S13. Whole-genome sequencing coverage breadth confirms loss of chromosome 3 in *Bso* MF6 mutant clones. Related to Fig. 3.** Per-clone mapping statistics (nanopore and Illumina reads aligned to the MF6 reference assembly) for each of the three *Bso* MF6 chromosomes and the mini-Tn5 transposon, including number of mapped reads, covered bases, percent breadth of coverage, mean depth, and mean mapping quality (MAPQ ≥ 10). Clones with chromosome 3 breadth of coverage near 0% (e.g., 20D2, 24G11) confirm complete loss of chromosome 3, consistent with the asterisks marking these strains in Fig. 3a.

**Supplementary Table S14. Candidate toxin proteins in the *Bso* MF6 pellet proteome. Related to Fig. 3.** Mean core-centered $log_2$ LFQ intensity per strain × condition group for the 34 candidate toxin shown in Fig. 3d. Values are given for wild-type *Bso* MF6, 27D6, and 34F7, grown alone (**−**) or in co-culture with Chlamydomonas (**+**) at day 2. Blank cells indicate the protein was not detected in that group; for the seven chromosome 3-encoded proteins, all 27D6 and 34F7 columns are blank because both mutants have lost chromosome 3. DeepLocPro_localisation, toxin_Pfam_domains, and T6SS_effector_probability are as described for Supplementary Table S10.

**Supplementary Table S15. Log2 MaxLFQ intensities per protein per proteomic run, culture supernatant of *Bso* MF6, 27D6 and 34F7. Related to Fig. 3 and Extended Data Fig. 4.** Log2-transformed MaxLFQ intensities, with the cross-run normalisation factor removed, for each bacterial protein detected in the culture supernatant (Sup) at day 2 across all 24 supernatant proteomic runs (genotype [27D6/34F7/MF6] × condition [C−, alone; C+, with Chlamydomonas] × biological replicate [1–4]; column headers follow the format genotype_condition_sup_replicate). One row per protein: Protein gives the locus tag; Description, chromosome, start, stop and strand give the annotation and genomic position; protein_length_aa gives the protein length; and n_peptides gives the number of peptides identified for the protein. Blank cells indicate the protein was not quantified in that run. The table has 378 entries:

377 individual proteins and one merged entry, GroEL_family, which combines three GroEL paralogs (AC1V0C_03900, AC1V0C_28400 and AC1V0C_34525). For the three chromosome 3-encoded proteins (AC1V0C_32450, AC1V0C_34715 and AC1V0C_34970), all 27D6 and 34F7 columns are blank because both mutants have lost chromosome 3.

**Supplementary Table S16. Differential protein abundance and presence/absence calls in the culture supernatant of *Bso* MF6, 27D6 and 34F7 with vs without Chlamydomonas. Related to Fig. 3 and Extended Data Fig. 4.** Long-format results for the supernatant (Sup) proteome (DIA), one row per protein per contrast, for the three co-culture versus monoculture comparisons (contrast column; group_A vs. group_B): MF6_C+_sup vs. MF6_C−_sup, 27D6_C+_sup vs. 27D6_C−_sup and 34F7_C+_sup vs. 34F7_C−_sup. logFC is group_A minus group_B, with accompanying p_value and Benjamini–Hochberg-adjusted p_adj [limma moderated t-test, as in Supplementary Table S10]; n_detected_A and n_detected_B give the number of replicates (of four) in which the protein was quantified in each group. Proteins detected in at least three replicates of one group and in none of the other [and passing an intensity criterion, define] are called ON/OFF instead of being tested: ON_group gives the group in which the protein is treated as ON and ON_percentile gives its LFQ percentile in the opposite group (as in Supplementary Table S10), and logFC, p_value and p_adj are blank for these rows (53 calls: 50 ON with *Chlamydomonas* and 3 ON in monoculture). UP/ON in MF6 only is set to 1 for the 35 proteins that are significantly more abundant (log2FC > 1, p_adj < 0.05) or ON in MF6 co-culture, but not in either mutant [confirm rule].

DeepLocPro_localisation, toxin_Pfam_domains, T6SS_effector_probability and the antiSMASH/MIBiG columns are as described for Supplementary Table S10. The table has 616 rows covering 297 proteins (194 rows for MF6, 181 for 27D6, 241 for 34F7), with p_adj < 0.05 for 35, 48 and 43 proteins, respectively. The three chromosome 3-encoded proteins are present only in the MF6 contrast because both mutants have lost chromosome 3. The merged GroEL_family entry is as in Supplementary Table S15.

**Supplementary Table S17. Primary root elongation of Arabidopsis seedlings mono-associated with *Bso* MF6 transposon mutants. Related to Fig. 4.** Per-seedling primary root elongation (cm) of Arabidopsis seedlings mono-associated with wild-type *Bso* MF6, each of the 14 non-inhibitory transposon mutants, or $MgCl_2$ buffer alone (control), across five independent experiments (Exp: mono_1–mono_4, mono_6). Strain gives the reported strain name; Strain_original and Tn give the as-isolated clone code and original Tn5 insertion identifier prior to consolidation (see Supplementary Table S12). Mutants represented by more than one independently isolated clone (27D6, 50A5) pool data from all isolates under the reported strain name.

**Supplementary Table S18. Leaf area of Arabidopsis seedlings mono-associated with *Bso* MF6 transposon mutants. Related to Fig. 4.** Per-seedling leaf area ($cm^2$) of Arabidopsis seedlings mono-associated with wild-type *Bso* MF6, each of the 14 non-inhibitory transposon mutants, or $MgCl_2$ buffer alone (control), across five independent experiments (Experiment: mono1–mono4, mono6). strain gives the reported strain name; strain_original and Tn give the as-isolated clone code and original Tn5 insertion identifier prior to consolidation (see Supplementary Table S12). Mutants represented by more than one independently isolated clone (27D6, 50A5) pool data from all isolates under the reported strain name. rep identifies the biological replicate within each strain/experiment combination.

**Supplementary Table S19. Differential gene expression at each timepoint relative to uninfected Chlamydomonas (T0). Related to Fig. 5.** DESeq2 differential expression results for 14,732 genes at each of four infection timepoints versus T0 (comparison: T1–T4_vs_T0), with $\log_2$ fold-change, standard error, *p*-value, and Benjamini-Hochberg adjusted *p*-value (*padj*). Membership_Score and Cluster_name give the fuzzy c-means membership score and assigned temporal expression cluster (Up 1–6, Down 7–12; see Fig. 5a, b) for dynamically expressed genes. geneSymbol, Description, Predalgo (predicted subcellular localization), GO, and Best-hit-arabi-name (closest Arabidopsis ortholog) provide functional annotation.

**Supplementary Table S20. ImpulseDE2 dynamic expression classification during Chlamydomonas infection. Related to Fig. 5.** ImpulseDE2 output testing each of 14,732 genes for dynamic expression across infection (single-condition mode). The 6,817 dynamically expressed genes (category ≠ NA) form the input to the temporal clustering shown in Supplementary Table S19 and Fig. 5a, b. geneSymbol, Description, Predalgo, GO, and Best-hit-arabi-name are as in Supplementary Table S19.

**Supplementary Table S21. GO term enrichment for the twelve temporal expression clusters. Related to Fig. 5 and Extended Data Fig. 5.** topGO enrichment results (weight01 algorithm, Fisher's exact test) for Gene Ontology Biological Process (BP), Cellular Component (CC), and Molecular Function (MF) terms within each of the twelve temporal expression clusters (Up 1–6, Down 7–12; see Supplementary Table S19, Fig. 5a, b). Significant and Annotated give the number of cluster genes and universe genes annotated with the term, respectively; Expected is the count expected by chance; weight01Fisher and neg_log10_p give the *p*-value and its −log10 transform. geneID lists the contributing genes (pipe-delimited).

**Supplementary Table S22. GO term enrichment for the merged acute-response gene set (Up 1 and Down 12). Related to Extended Data Fig. 5.** topGO enrichment results (weight01 algorithm, Fisher's exact test) for the merged gene set comprising temporal clusters Up 1 and Down 12. The acute, transiently induced/repressed clusters underlying the flagellar regeneration signature described in the main text and shown in Extended Data Fig. 5b. Columns as in Supplementary Table S21.

**Supplementary Table S23. Expression of a manually curated gene set across infection. Related to Extended Data Fig. 5.** $Log_2$ fold-change and DESeq2 adjusted *p*-value at each infection timepoint (T1–T4) relative to T0, for a manually curated list of genes shown in Extended Data Fig. 5c: representative genes from each temporal expression cluster, together with additional genes of interest. Cluster gives the assigned temporal expression cluster (Up 1–6, Down 7–12; see Supplementary Table S19) where applicable; blank for not temporally expressed genes. Alias gives the common gene name where available. Description, Predalgo, GO, and Best-hit-arabi-name are as in Supplementary Table S19.

**Supplementary Table S24. Expression of candidate programmed cell death (PCD)-related genes across infection. Related to Extended Data Fig. 5.** $Log_2$ fold-change and DESeq2 adjusted *p*-value at each infection timepoint (T1–T4) relative to T0, for 19 candidate PCD-related genes selected based on homology to conserved algal cell-death genes identified in Mizrachi et al.[5], shown in Extended Data Fig. 5d. Cluster gives the assigned temporal expression cluster (Up 1–6, Down 7–12; see Supplementary Table S19) where applicable; blank for not temporally expressed genes. Alias, Description, Predalgo, GO, and Best-hit-arabi-name are as in Supplementary Table S19.

**Supplementary Table S25. Expression of autophagy (ATG) machinery and TOR-pathway genes across infection. Related to Extended Data Fig. 5.** $Log_2$ fold-change and DESeq2 adjusted *p*-value at each infection timepoint (T1–T4) relative to T0, for 24 core autophagy (ATG) and TOR-pathway genes shown in Extended Data Fig. 5e. Cluster gives the assigned temporal expression cluster (Up 1–6, Down 7–12; see Supplementary Table S19) where applicable; blank for not temporally expressed genes. Alias, Description, Predalgo, GO, and Best-hit-arabi-name are as in Supplementary Table S19.

**Supplementary Table S26. Expression of zinc/iron transporter and metal tolerance genes across infection. Related to Fig. 6.** $Log_2$ fold-change and DESeq2 adjusted *p*-value at each infection timepoint (T1–T4) relative to T0, for 16 ZIP/ZRT/IRT/MTP-family metal transporter and tolerance genes shown in Fig. 6e, including ZIP3 (Cre03.g189550). Cluster gives the assigned temporal expression cluster (Up 1–6, Down 7–12; see Supplementary Table S19) where applicable; blank for not temporally expressed genes. Alias, Description, Predalgo, GO, and Best-hit-arabi-name are as in Supplementary Table S19.
